# Endocrine disruption of early uterine differentiation causes adenocarcinoma mediated by Wnt/β-catenin- and PI3K/AKT signaling

**DOI:** 10.1101/2022.11.18.517135

**Authors:** Elizabeth Padilla-Banks, Wendy N. Jefferson, Brian N. Papas, Alisa A. Suen, Xin Xu, Diana V. Carreon, Cynthia J. Willson, Erin M. Quist, Carmen J. Williams

**Affiliations:** Reproductive and Developmental Biology Laboratory, National Institute of Environmental Health Sciences, National Institutes of Health, Research Triangle Park, NC 27709, USA; Epigenetics and Stem Cell Biology Laboratory, National Institute of Environmental Health Sciences, National Institutes of Health, Research Triangle Park, NC 27709, USA; Inotiv-RTP, Research Triangle Park, NC 27709, USA; Experimental Pathology Laboratories, Research Triangle Park, NC 27709, USA

**Author notes:** These authors contributed equally to the manuscript (Elizabeth Padilla-Banks, Wendy N. Jefferson). Present address: Charles River Laboratories, Durham, NC 27703, USA. **Correspondence:** Carmen J. Williams.

**Keywords:** Endometrium, epithelial stem cells, scRNAseq, spatial sequencing, adenocarcinoma, diethylstilbestrol

## Abstract

Developmental exposure to non-mutagenic environmental factors can contribute to cancer risk, but the underlying mechanisms are not understood. We used a mouse model of endometrial adenocarcinoma that results from brief developmental exposure to an estrogenic chemical, diethylstilbestrol (DES), to determine causative factors. Single cell RNA sequencing and spatial transcriptomics of adult uteri revealed new markers of uterine epithelial stem cells, identified luminal and glandular progenitor cells, and defined a glandular epithelial differentiation trajectory. Neonatal DES exposure disrupted uterine epithelial differentiation, resulting in widespread activation of Wnt signaling and a failure to generate epithelial stem cells or distinguishable glandular and luminal epithelial cells. The endometrial stromal cells activated inflammatory signals and oxidative stress. Together, these changes activated PI3K/AKT signaling to drive malignant transformation. These findings explain how human cancers, which are often associated with abnormal activation of PI3K/AKT signaling, could result from exposure to environmental insults during development.

**Highlights:**

- Single cell analysis of adult mouse uteri reveals epithelial stem cell markers and gland development trajectory
- Developmental DES exposure causes widespread activation of Wnt signaling and failure of epithelial cell differentiation
- Uterine adenocarcinoma results from combined Wnt/β-catenin activation, stromal inflammation, and induction of PI3K/AKT signaling
- OLFM4 is a marker of developmental DES-induced uterine adenocarcinoma

## Introduction

Cellular differentiation is genetically and epigenetically programmed but strongly influenced by the environment. During development, the influence of environmental factors is integrated by diverse cues that come from within the organism, the maternal environment, and the external environment. Environmental cues provide opportunities for the developing organism or differentiating adult tissues to adapt their pre-programmed differentiation trajectory to improve fitness^1,2^. Altered developmental trajectories can also reduce adult fitness, particularly in settings where the environment changes after there are developmental adaptations to the initial environment. This phenomenon, now termed “developmental origins of health and disease (DOHAD)”, was first discovered as a connection between poor nutrition in infants and increased risk of arteriosclerotic heart disease in prosperous adults^3,4^.

Environmental exposures during specific windows of development can even lead to delayed onset of cancer in adulthood. A classic example of this phenomenon is prenatal exposure to diethylstilbestrol (DES), which alters developmental patterning of the female reproductive tract and leads to an increased incidence of specific cancers in adult women^5–7^. The “two-hit hypothesis” of cancer development postulates that two mutational events cause cancer, with the first mutation causing cancer susceptibility and the second mutation causing cancer progression^8–10^. Indeed, numerous cancers result from two mutational events, such as retinoblastoma and colorectal carcinoma^11,12^. In addition to mutations, epigenetic alterations can similarly serve as “hits” responsible for cancer development through their impacts on tumor suppressors or oncogenes^13^.

But how do non-mutagenic environmental exposures during development impact tissues in a way that leads to late onset of carcinogenesis, particularly when similar exposures during adulthood have no persistent effects? To address this question, we are utilizing a mouse model of DES exposure during female reproductive tract differentiation^14^. At birth, the mouse female reproductive tract is a bifurcated tubular structure lined by a simple epithelium. The uterus is not fully differentiated until about three weeks later, when anterior-posterior patterning and cellular differentiation are fully established under the influence of factors including *Hox* and *Wnt* genes, growth factors, Hippo signaling and steroid hormones^15,16^. The mature uterine endometrium has a luminal epithelium and a glandular epithelium formed from invaginations of the luminal epithelium into the underlying stroma. These epithelia undergo cyclic regeneration with each estrous cycle from rare adult epithelial stem cells, which provide progenitor cells committed to proliferate and differentiate into luminal or glandular epithelium^17,18^.

The mouse model of neonatal DES exposure entails exposing newborn female mice to DES daily for 5 days beginning on the day of birth, during the initial window of rapid reproductive tract differentiation. This exposure causes altered reproductive tract patterning, increased deposition of extracellular matrix material throughout the tissue, and a high incidence of estrogen receptor alpha (ERα)-dependent uterine endometrioid adenocarcinoma in 12 month old adults^14,19,20^. This model requires two “hits”: neonatal exposure to an estrogenic chemical followed by additional exposure to endogenous estrogen following puberty. The resulting cancers have no obvious mutational signature and are not discrete tumors^14,21–23^. Instead, they are characterized by having multiple foci of diffuse infiltrating neoplastic cells with mixed glandular, basal and squamous cell features^14,20^.

To determine the mechanisms underlying neonatal DES exposure-induced cancer development we utilized single cell RNA sequencing (scRNAseq) and spatial transcriptomics^24^ to identify alterations in uterine gene expression in adult DES-exposed mice with adenocarcinoma relative to unexposed controls. We found that DES-exposed mice lack normally differentiated uterine luminal and glandular epithelial cells. Instead, they have an expanded population of less differentiated uterine epithelial cells that have characteristics of stem and progenitor cells and extensive activation of Wnt/β-catenin signaling. The endometrial stroma is highly enriched in inflammatory pathways, and the cancer cells have activated PI3K/AKT signaling, a common driver of uterine adenocarcinoma. These findings indicate that brief estrogenic chemical disruption of uterine epithelial cell differentiation, combined with stromal inflammation, promotes abnormal activation of Wnt/β-catenin and PI3K/AKT signaling pathways that drive cancer development.

## Results

### Identification of uterine cell types in control and DES-exposed mice

Single cells were isolated from a single sample of pooled uteri of control (CO) or DES-exposed mice, processed using the 10x Genomics scRNAseq platform, and sequenced to a depth of >1.2 billion reads per sample with >40,000 reads per cell. Mapping was of high quality and 32387 CO cells and 16312 DES cells had sufficient information captured (Table S1A). There was a robust number of transcripts per cell (CO, 6779; DES, 10153). Unsupervised clustering of all cells was performed using Seurat v3.1.0, and a Uniform Manifold Approximation and Projection (UMAP) generated (Figure 1A). Clustered cell types were identified by comparing differentially expressed genes (DEGs) of each cluster to published uterine cell type markers^25^ (Figure 1B; Table S1B). The largest clusters were epithelial and mesothelial cells, but there was representation of most expected cell types. Each treatment group also had unique cell type clusters (Figure 1B). One CO cluster strongly expressed the secreted factor *Ctla2a* (cytotoxic T lymphocyte-associated protein 2 alpha). A small CO cluster of apparent mesothelial cells overexpressed the microtubule regulator *Stmn1* (stathmin 1). Unique to the DES sample, a cluster of myeloid cells overexpressed *Lyz2* (lysozyme 2). Three cell types found in the uterine stromal region were largely absent from the DES samples: stromal cells, pericytes and endothelial cells.

**Figure 1:**
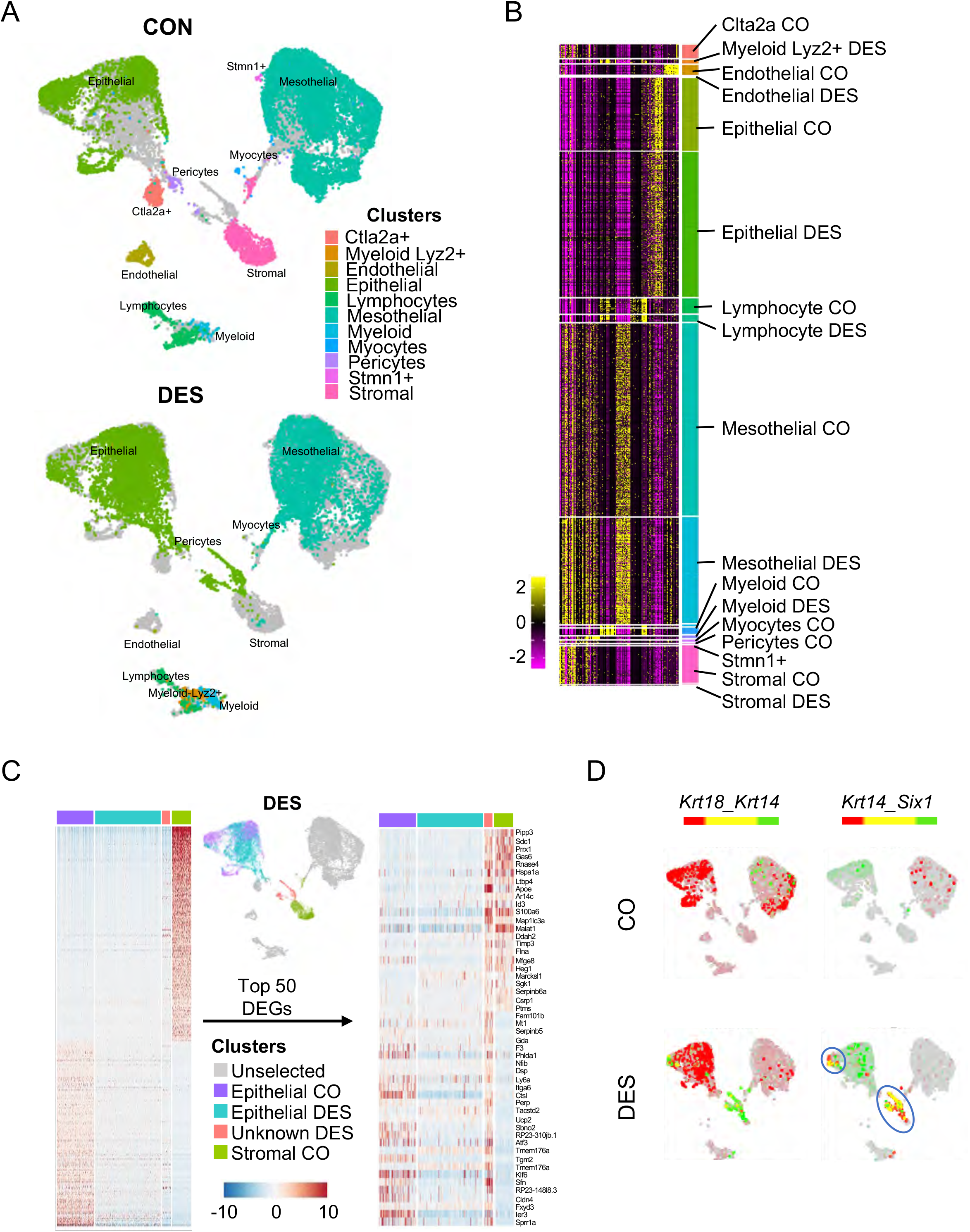
Identification of uterine cell types captured by scRNAseq. A) Integrated UMAP of all uterine cells captured. Each cell type identified by color and text from CO (top) and DES (bottom). B) Heat map of gene expression of cell identity genes. Cell type indicated for each identified cluster. Expression is centered [mean=0 ± standard deviation (SD) of each feature]. C) Heat map of epithelial and stromal cell markers in CO and DES epithelial cells, CO stromal cells and an unidentified population with similarities to both cell types (left). Top 25 cell identity DEGs are plotted for epithelial and stromal cells. Expression is Pearson Residuals from the SCTransform method. D) Dual feature plots of epithelial and basal cell markers (*Krt18, Krt14* and *Six1*). CO (top) and DES (bottom). In this and all subsequent dual feature plots, colors for each gene (red or green) are indicated above the UMAPs; yellow indicates overlapping expression.

The lack of stromal cells in the DES sample was surprising compared to the robust capture of CO stromal cells. One explanation for this finding was that another cell type in the DES sample was not categorized properly. Indeed, there was a group of DES cells that categorized as epithelial cells but plotted closer on the UMAP to CO stromal cells and mesothelial cells than to CO epithelial cells. To test for mis-categorization, we first performed unbiased clustering of DES epithelial cell gene expression with CO mesothelial and stromal cell gene expression. There was no resulting subset of DES mesothelial cells that had characteristics of stromal cells (Figure S1A). Next, a comparison of DES epithelial cell gene expression to both CO epithelial and stromal cells revealed a subcluster of DES cells that exhibited characteristics of both cell types (Figure 1C). This finding was confirmed by a heat map of the top 50 DEGs differentiating CO epithelial from CO stromal cells (Figure 1C). DEGs in this DES subcluster included the basal cell markers *Trp63* and *Krt14*. Neonatal DES exposure causes development of a population of basal cells, not observed in controls, that co-express *Trp63, Krt14*, and *Six1* ^20,26,27^. Feature plots of the luminal epithelial cell marker, *Krt18*, and these basal cell markers clearly identified two DES cell clusters as basal cells. One of these clusters was near the CO stromal cells and one was adjacent to the main epithelial cell cluster; this cell type was not observed in CO cells (Figures 1D and S1B). These findings confirm that basal cells were included in the analyzed DES cell populations. Feature plots of three stromal cell markers, *Dpt*, *Vcan* and *Col6a3*^28^, confirmed that the DES group lacked stromal cells (Figure S1C). It is likely that increased levels of extracellular matrix deposition in the stroma of DES uteri precluded isolation of living stromal cell types for analysis^14,29^. *Dpt* and *Vcan* were highly expressed in the CO *Ctla2+* cells, suggesting them as a stromal cell population (Figure S1C).

### Comparison of CO and DES stromal cells using spatial transcriptomics

Because DES stromal cells were not captured in our scRNAseq analysis, we used spatial transcriptomics to characterize gene expression in DES stromal cells. Frozen longitudinal sections from uteri of 12-month-old CO and DES mice were evaluated to ensure inclusion of the luminal region. DES sections were also evaluated to determine the presence of uterine adenocarcinoma. Once appropriate regions were confirmed in two tissue blocks per group, immediately adjacent sections were processed for spatial transcriptomics (Figure 2A). There were ~40 million total reads per sample. One of the CO sections had a substantial tissue fold and could not be analyzed further and one of the DES sections was limited in identifying clusters so only one section per group was chosen for further analysis (Figure S2A). The CO sample had 1433 spots under the tissue with 56% sequencing saturation (28910 mean reads per spot) and a median of 2581 genes per spot. The DES sample had 1575 spots under the tissue with 65% sequencing saturation (25666 reads per spot) and a median of 2056 genes per spot. Note that each spot, which is 55 μm in diameter (10x Genomics), represents gene expression from approximately 5-10 cells, depending on cell size. For this reason, clear distinctions in gene expression cannot be made for intermixed cell types using this methodology.

**Figure 2:**
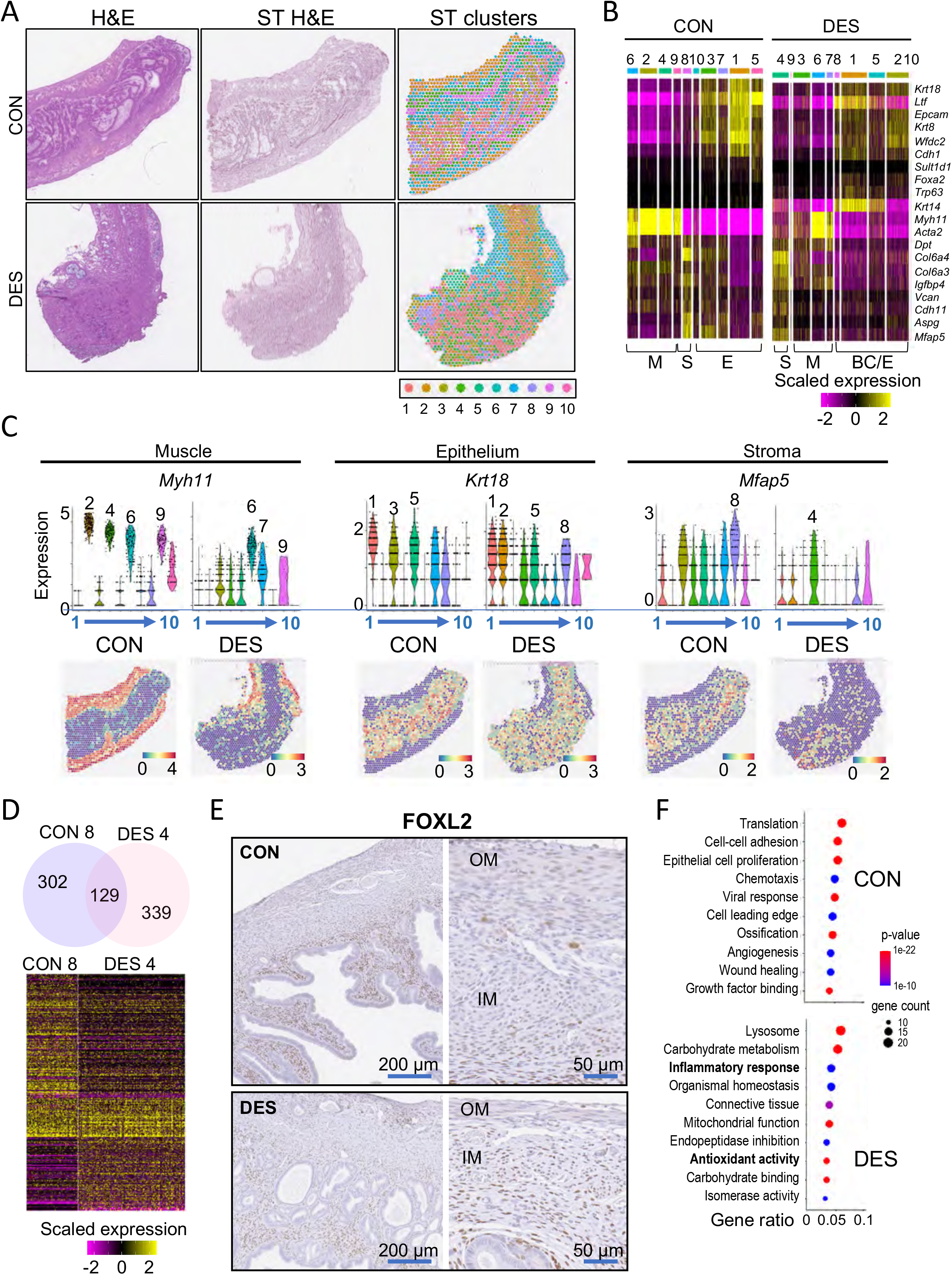
Stromal cell gene expression is altered by DES as evidenced by spatial transcriptomics. A) Uterine tissue sections from CO and DES used for spatial transcriptomics (ST). H&E stain of tissue section adjacent to ST section (left) and tissue sections used for ST (ST H&E, middle). Cluster identification using Space Ranger-1.2.2, skmeans 10 (10x Genomics); colors represent distinct clusters (right). B) Heat map of select uterine tissue cell type markers plotted for CO (left) and DES (right). Cell type is indicated below heat maps (M=muscle, S=stroma, E=epithelium, BC=basal cells). Expression values are Pearson Residuals from the SCTransform method. C) Violin plots and ST section of select markers for CO and DES (cluster number indicated below plot). Violin plot expression same as panel B; ST expression (natural log transformed counts). D) Venn diagram of CO and DES stromal cluster specific DEGs (top). Heat map of CON and DES stromal cell markers (bottom). Expression same as panel B. E) Immunohistochemistry of FOXL2 in 12-month-old CO and DES exposed uteri. CO (top) and DES (bottom); right panels are higher magnification of each. OM, outer muscle; IM, inner muscle. F) Top 10 non-overlapped GO categories for CO (top) and DES (bottom). Gene ratio (#genes in GO category/#DEGs), p-value and gene count indicated.

Spot clusters were distinguished in both CO and DES tissue sections using Space Ranger-1.2.2, skmeans 10 (10x Genomics) (Figure 2A; Tables S2A-B). Cluster cell type identification was made using published markers (Figure 2B)^20,25,28,30,31^. CO sections had clusters identifiable as regions of muscle, stroma, and epithelium (Figures 2A and 2B). Glandular epithelium was present in CO cluster 5 based on having high expression of *Foxa2* and *Sult1d1*. CO cluster 10 could not be characterized as one specific cell type and likely represented a mixture of epithelial and stromal cells in close proximity. The DES clusters had regions of muscle, stroma, and regions representing a mixture of epithelial and basal cells (Figures 2A and 2B). Cell type identification was confirmed using violin plots of a highly expressed marker for each cell type combined with spatial expression plots showing their histologic locations (Figure 2C). Stromal cells were identified clearly in DES cluster 4.

To explore gene expression differences between CO and DES stromal cells, we overlapped the 431 DEGs in CO cluster 8 (Table S2A) with 468 DEGs in DES cluster 4 (Table S2B). Only about 30% of the DEGs were in common, including stromal cell markers *Vcan, Dpt* and *Col6a3* (Figure 2D, Tables S3A, S3B and S3C). Interestingly, the well characterized stromal cell marker, *Foxl2*^3^ was a DEG in CO but not DES stromal cell clusters. Immunohistochemical staining of FOXL2 in CO sections revealed strong staining in the inner stroma adjacent to epithelium and weaker staining in the outer stroma and muscle layers (Figures 2E and S2B). In DES uterus sections, FOXL2 staining was similar across stroma and muscle, explaining why *Foxl2* was not a DEG for DES stroma.

The most significantly enriched biological processes identified by GO analysis of CO-specific stromal DEGs included extracellular matrix organization, translation, and regulation of signal transduction (Table S3D). GO analysis of DES-specific stromal DEGs also identified extracellular matrix organization, but unlike the GO categories for CO DEGs, oxidative phosphorylation and glycolytic process were identified as significantly enriched (Table S5E). The top non-overlapping GO categories revealed substantial differences between CO and DES (Figure 2F; Tables S5F and S5G). Relevant to cancer development, DES stromal cells were particularly enriched in categories related to antioxidant activity and inflammatory responses.

### DES epithelial cells lack luminal or glandular identity

To determine how epithelial cell types differed between CO and DES samples, we restricted our scRNAseq analysis to include only cells differentially expressing the epithelial cell marker *Krt18*. Integrated UMAP analysis identified four distinct cluster regions with almost no overlap between CO and DES cells (Figure 3A). To determine which cell types were present in each cluster, marker genes for each cluster were determined by Seurat+MAST (Fig S3A; Table S4). Two DES-specific clusters (8 and 11) separate from the main grouping of DES cells differentially expressed two basal cell markers (*Krt14* and *Trp63*) and *Six1*, which is mainly expressed in basal cells (Figure 3A; Table S4)^20^. Dual feature plots confirmed the identity of these two DES clusters as basal cells based on their high, overlapping expression of *Krt14* and *Trp63* (Figure 3A). As expected, CO epithelial cell clusters lacked the basal cell markers.

**Figure 3:**
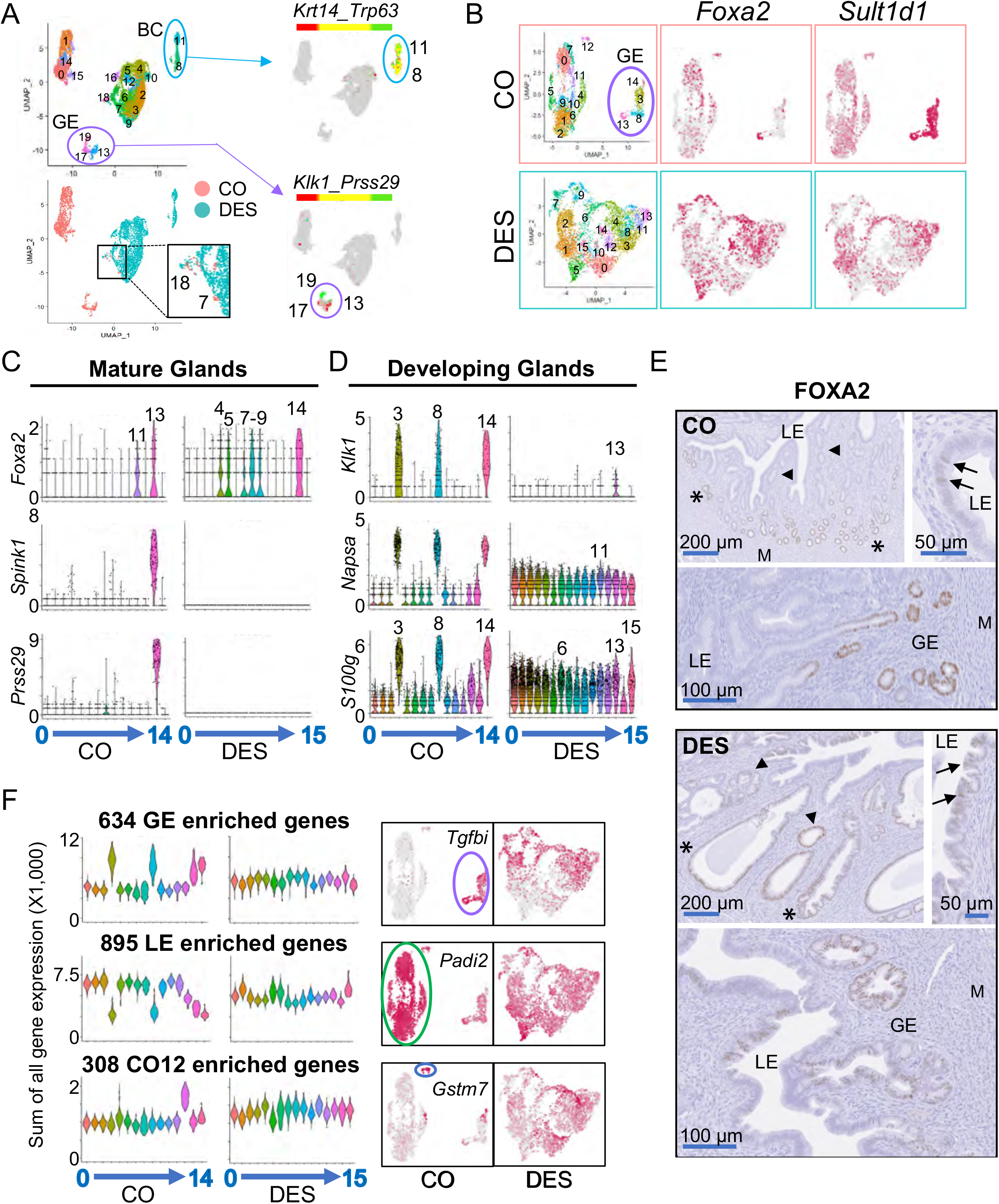
Epithelial cells from DES exposed mice lack subtype identity. A) UMAP of integrated epithelial cells from CO and DES scRNA-seq samples (top left); cluster numbers indicated. Epithelial cells identified as either CO or DES on the integrated UMAP (bottom left). Overlapping CO and DES cells found in only one region (clusters 7 and 18, magnified area indicated). Dual feature plots (right) for basal cell (BC) markers (*Krt14* and *Trp63*) and glandular epithelial (GE) markers (*Klk1* and *Prss29*). B) UMAPs of non-integrated CON and DES epithelial cells. GE clusters outlined in purple. Feature plots of *Foxa2* and *Sult1d1* for CO (top) and DES (bottom). C) Violin plots of mature GE markers for CO and DES; cluster numbers on x-axes. Expression levels indicated on y-axes as natural log transformed counts. High expression cluster numbers indicated. D) Violin plots of developing GE markers for CO and DES; cluster numbers on x-axes. Expression levels indicated on y-axes as natural log transformed counts. High expression cluster numbers indicated. E) Immunohistochemistry of FOXA2 in CO (top) and DES (bottom). LE, luminal epithelium; GE, glandular epithelium; M, myometrium; asterisks, outer GE; arrowheads, inner GE; arrows, LE FOXA2 expression. F) Violin plots of summed expression (SCTransform counts X 1,000) of GE, LE and CO cluster 12 (CO12) specific DEGs for CO and DES epithelial cell clusters. Cluster numbers on x-axes. Feature plots of a representative gene from each category (right).

A group of three CO-specific clusters (13, 17, and 19) were separate from the largest grouping of CO cells (Figures 3A and S2A, Table S6). These clusters differentially expressed several uterine gland-specific genes, including *Prss29, Spink1, Sult1d1, Napsa, Gpx3* and *Klk1*, indicating that they represented glandular epithelium (GE) (Figure S3A, Table S4)^16,32^. Of these markers, *Prss29* and *Spink1* are highly expressed in mature GE whereas *Napsa* and *Klk1* are expressed in developing GE. Dual feature plots of *Klk1* and *Prss29* showed non-overlapping expression in these clusters with *Klk1* expressed in clusters 13 and 17 and *Prss29* expression in cluster 19, suggesting clusters 13 and 17 were developing GE and cluster 19 was mature GE (Figure 3A). The large grouping of CO epithelial cell clusters (0, 1, 14, 15) were presumed luminal epithelial (LE) cells as they comprised the largest number of epithelial cells in the mouse uterus (Figure 3A). The large grouping of DES epithelial cells did not overlap with either the presumed CO LE cells or the CO GE though there were a few CO cells clustered with this DES group (Figures 3A and S3A, Table S4).

The lack of overlap in UMAP locations of the CO and DES epithelial cells indicated that these populations were quite different in their gene expression and precluded further joint analysis of subclusters. Instead, we used a non-integrated approach to identify subclusters in CO and DES epithelial cells independently. We first removed the basal cells from the DES epithelial cells based on their expression of both *Krt14>1* count and *Trp63>1* count and then performed a separate UMAP analysis of the remaining epithelial cells in each group. This analysis identified 15 CO clusters and 16 DES clusters (Figure 3B; Tables S5A-B). The CO clusters contained 2629 cells with three distinct groups of cells: a small single cluster 12, a group containing clusters 14, 3, 8 and 13 and a large group containing all other clusters. The DES epithelial clusters contained 4610 cells that were not clearly separated into groups, indicating that they had less differential gene expression between clusters. Feature plots of the mature GE marker, *Foxa2*, showed strong expression in CO cluster 13, whereas the general GE marker, *Sult1d1*, was strongly expressed in CO clusters 3, 8, 13 and 14, indicating that this group of 4 CO clusters was the GE cells (Figure 3B). In the DES cells, feature plots of *Foxa2* and *Sult1d1* showed enriched expression of both in multiple clusters but no distinctly identifiable GE clusters (Figure 3B).

To further confirm the identity of the putative CO GE clusters and to identify GE in DES clusters, violin plots were generated for genes expressed in mature GE cells including *Foxa2, Spink1*, and *Prss29*^32^ (Figure 3C). CO cluster 13 highly expressed all three markers, indicating that this cluster contained the mature GE cells. *Foxa2* was also expressed in CO cluster 11 within the putative LE cell grouping; this cluster lacked expression of *Spink1* and *Prss29*, indicating that these cells were not mature GE. In the DES cells, *Foxa2* was highly expressed in multiple DES clusters but there was no corresponding expression of *Spink1* or *Prss29*, indicating that none of these clusters were mature GE. The close proximity of CO clusters 3, 8 and 14 to the *Foxa2+* mature GE CO cluster 13 suggested that these clusters might represent GE in an earlier stage of differentiation. To test this idea, we generated violin plots of developing GE markers^16^ (Figure 3D). High expression of *Klk1* and *Napsa* in CO clusters 3, 8 and 14, but not 13, confirmed these clusters as developing GE. Several other genes exhibited the same expression pattern, including *S100g, Hif1a, Pim3, Prap1* and *Wnt7b* (Figures 3D and S3B). In the DES clusters, these markers were quite evenly distributed among almost all clusters, precluding identification of developing GE clusters (Figure 3D and S3B).

To confirm the identification of CO clusters as developing and mature GE and LE, we examined FOXA2 localization in uterine sections (Figures 3E and S3C). In CO uteri, FOXA2 was primarily found in GE; however, not all GE expressed FOXA2. FOXA2 staining was highest in the distal GE (farthest away from LE) and was not in invaginating GE or GE nearest the LE. Some luminal cells also expressed FOXA2, likely representing CO cluster 11 and supporting the identification of the large grouping of CO clusters as LE (Figures 3C, 3E and S3C). In DES uteri, FOXA2 was sporadically expressed in GE with no distinct pattern with respect to proximity to LE and was expressed in some areas of invaginating epithelium. There were many more FOXA2+ LE cells compared to the number in CO uteri. These findings indicated that some *Foxa2+* DES clusters were GE but that they did not exhibit the normal GE gene expression patterns observed in controls.

We next took a global approach to distinguish LE and GE specific gene expression in the DES clusters. The CO clusters were merged into three groups: 1. CO GE clusters (3, 8, 13 and 14); 2. CO LE clusters (0, 1,2, 4, 5, 6, 7, 9, 10 and 11); and 3. CO cluster 12. Genes that were increased 1.2-fold in one group over both other groups with padj<0.01 were considered “specific” to that group (Tables S6A-C). This procedure identified 634 GE-specific genes, 895 LE-specific genes and 308 CO cluster 12-specific genes. Violin plots of the sum of the gene expression in each category (GE, LE or CO12) observed in each individual CO cluster showed robust differential gene expression and easily identifiable patterns (Figure 3G). In DES clusters, this cell type-specific gene expression was not observed in summary violin plots of the cell type specific genes or the top five DEGs in each cell type category (Figures 3C, 3D, 3F and S3D). Feature plots of an example gene specific to each category confirmed the cell type-specific patterns in CO clusters; however, no specificity was observed in DES clusters (Figure 3F). These data demonstrate that DES epithelial cells lack cell type-specific gene expression.

### Differentiation trajectory of glandular epithelium in controls

The separation of CO GE into four clusters along with markers of mature GE in one cluster and developing GE markers in the other three suggested the GE clusters were at different stages of differentiation. To explore this idea further, we re-clustered only the four CO GE cell clusters followed by a slingshot analysis^33^ (Figure 4A). Ten resulting slingshot clusters had 1,205 up-regulated DEGs that represented only one trajectory pattern (Figure 4A; Table S7A). Progenitor cell markers *Foxm1* and *Top2a* were in slingshot cluster 6, developing GE markers *Klk1* and *Napsa* in slingshot clusters 0, 1 and 2, and mature GE markers *Foxa2, Spink1* and *Prss29* in slingshot clusters 5 and 7^16,32,34,35^. These markers confirmed the trajectory pattern and the direction of developmental gene expression.

**Figure 4:**
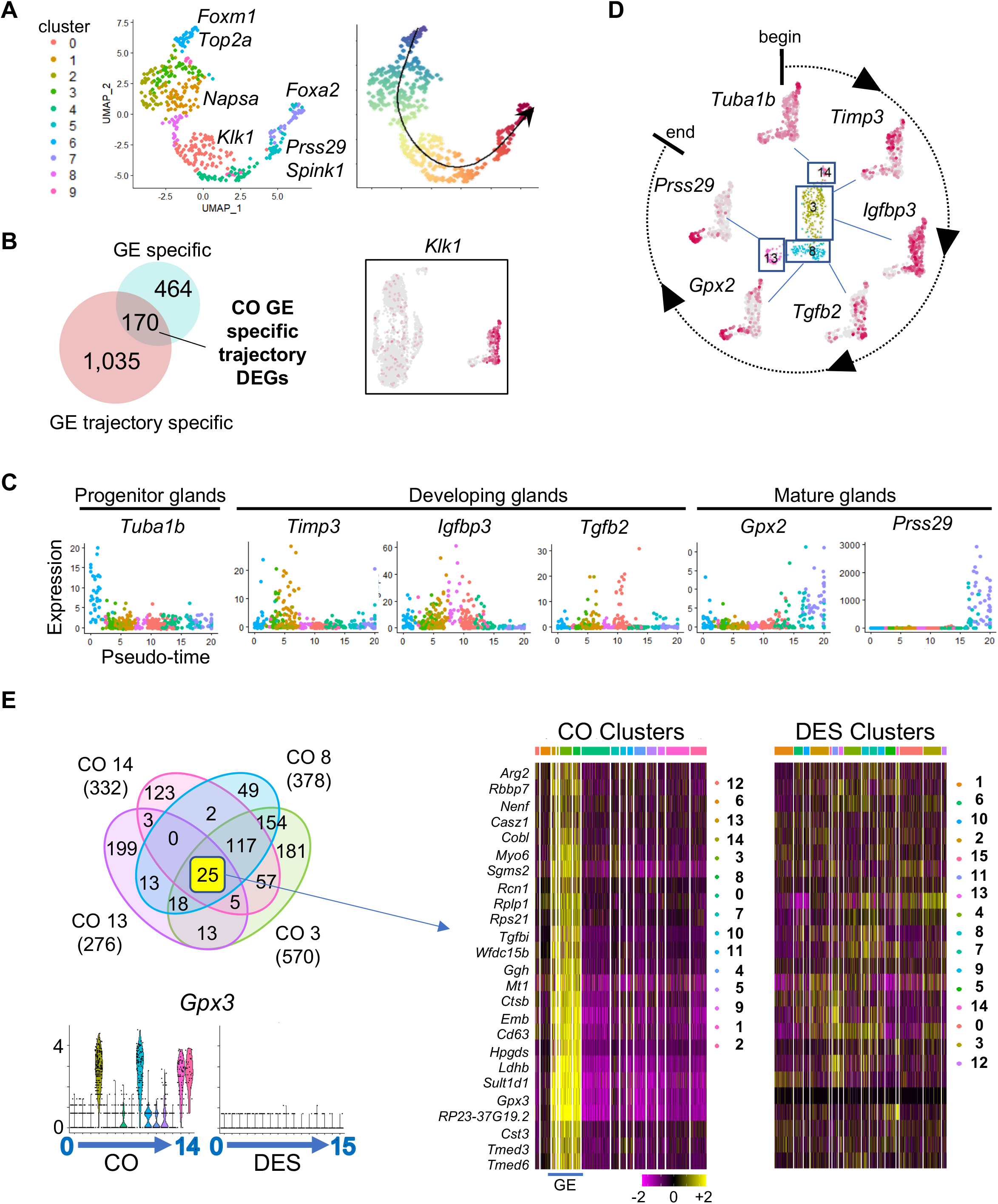
GE differentiation trajectory determined by Slingshot analysis is unidentifiable in neonatal DES exposed mice. A) UMAP of Slingshot analysis of CO GE cells (left). Cluster numbers designated by color; select genes indicated. Trajectory UMAP indicated by color (beginning, purple; end, red); arrow indicates direction of differentiation. B) Venn diagram of GE specific DEGs overlapped with GE trajectory specific DEGs. Feature plot of representative DEG from overlapped gene list, *Klk1*. C) Pseudo-time plots of select GE trajectory specific DEGs. Each dot represents one cell colored by cluster from panel A. D) Feature plots of CO GE using select genes from panel C; differentiation direction indicated. E) Venn diagram of DEGs from CO GE clusters 3, 8, 13 and 14 (top left). Violin plots of *Gpx3*, selected from 25 common DEGs (bottom left). Expression is natural log transformed counts. Hierarchical clustering heat map of 25 common DEGs in CO GE cells; cluster number indicated by color at top. DES clusters plotted using the same DEG order (right). Expression is centered (mean=0 ± SD of each feature).

To identify more precisely markers specific to different stages of GE development, we first restricted the slingshot cluster genes to GE-specific markers by excluding those also found in LE cell clusters, such as *Top2a* and *Foxa2*. Overlapping the 1,205 up-regulated slingshot cluster genes with 635 GE-specific up-regulated genes resulted in 170 GE-specific genes that contributed to the trajectory of GE cell development (Figure 4B; Table S7B). A feature plot of one of these genes, *Klk1*, demonstrated specificity to the developing GE clusters identified previously (Figure 4B). Pseudotime plots of six of the most highly differentially expressed GE-specific genes across the slingshot clusters demonstrated progression through differentiation (Figure 4C). This progression is summarized in a circle of differentiation time mapped back onto the non-integrated map of CO GE clusters (Figure 4D). These data confirmed that CO GE cluster 14 was the GE progenitor cell (GPC) population.

**Figure 5:**
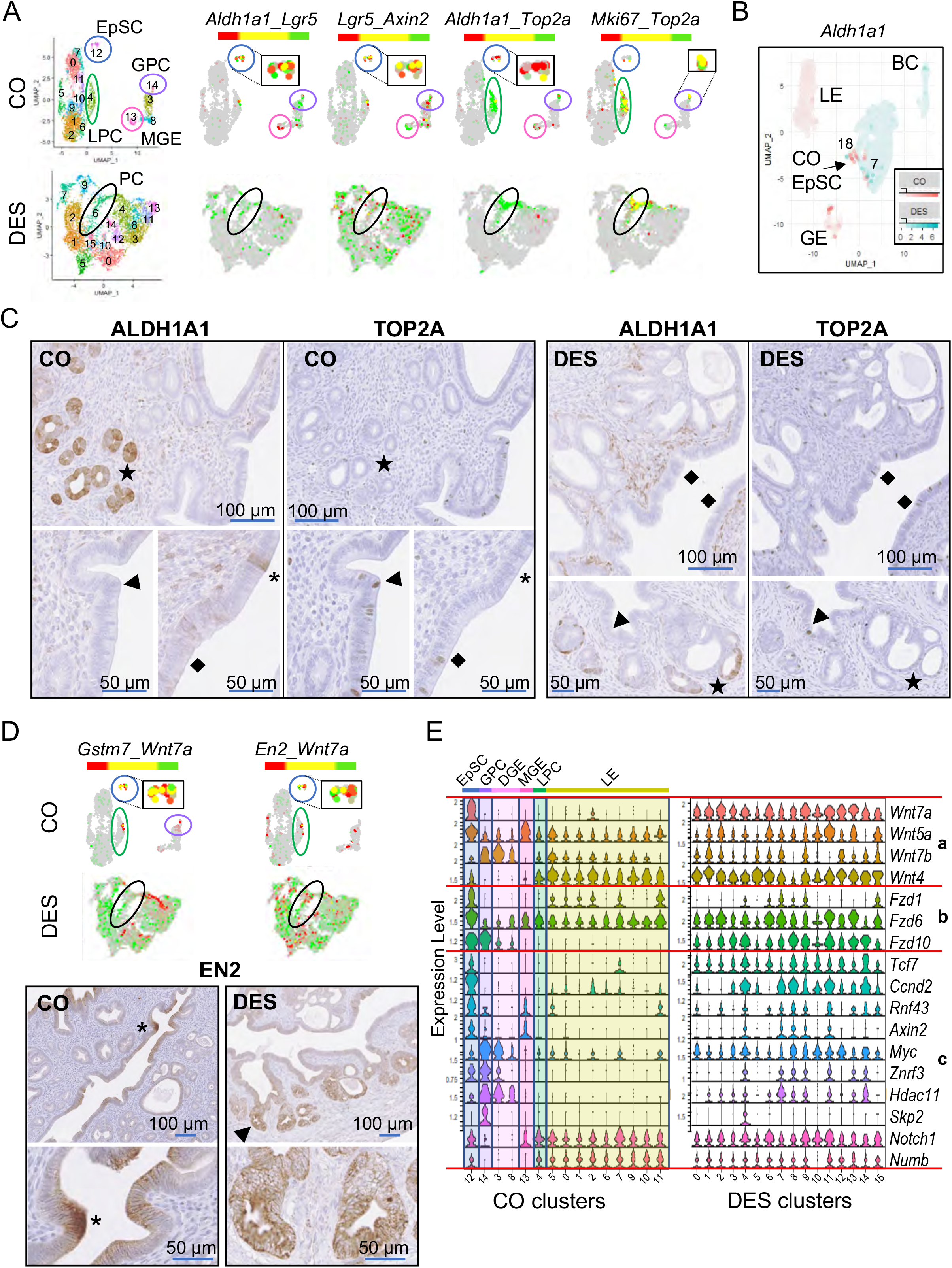
Epithelial stem cell and progenitor cell populations are substantially altered following neonatal DES exposure. A) UMAPs of non-integrated CON and DES epithelial cells. Epithelial cell populations identified as EpSC (epithelial stem cells, blue), GPC (GE progenitor cells, purple), LPC (LE progenitor cells, green), MGE (mature GE), and PC (progenitor cells, black). Dual feature plots of select stem cell (*Aldh1a1, Lgr5* and *Axin2*) and actively dividing cell (*Top2a* and *Mki67*) markers for CO (top) and DES (bottom). Insets in black boxes are EpSC or GPC magnified for clarity. B) Dual feature plot of *Aldh1a1* in CO and DES epithelial cells from integrated UMAP (CO, pink; DES, teal). Co EpSC, GE, LE and BC populations are indicated. C) Immunohistochemistry of ALDH1A1 and TOP2A in adjacent sections from CO and DES uteri, as indicated. Stars (GE) or asterisks (LE) indicate cells that are ALDH1A1+/TOP2A-. Arrowheads (GPC or LPC) and diamonds (LE) indicate cells that are ALDH1A1−/TOP2A+. Symbols correspond to same location in adjacent section. D) Dual feature plots of *Gstm7, En2* and *Wnt7a* for CO and DES. CO EpSC, GPC and LPC populations indicated as in panel A. Insets in black boxes are EpSC magnified for clarity. Immunohistochemistry of EN2 in CO (bottom left) and DES (bottom right) uteri. Asterisk, EN2 at invaginating GE; higher magnification of same region below. Arrowhead indicates region of high EN2 in disorganized GE; higher magnification of same region below. E) Violin plots of select Wnt ligands (a), receptors (b) and targets (c) in CO (left) and DES (right) epithelial clusters. Cluster numbers on x-axes; epithelial subtype in CO indicated at top. Expression is natural log transformed counts (y-axes).

*Foxa2* has long been used as a marker of GE; however, our scRNAseq data showed that *Foxa2* was expressed only in mature GE and in a small population of LE cells, findings that were validated at the protein level (Figure 3E). This observation is consistent with previously published localization of FOXA2 in adult uterus but differs from the ubiquitous presence of FOXA2 in the mature uterine glands of pregnant or pseudopregnant mice^36,37^. To find markers that would identify all developmental stages of GE but not LE, we selected the increased DEGs from all the CO GE clusters and overlapped them with each other (Figure 4E). Twenty-five DEGs were GE specific and found in all four CO GE clusters; a heat map confirmed their high exclusive expression in these clusters. For comparison, a heat map of the DES clusters for these 25 genes showed only sporadic expression in most clusters. Violin plots of one of these genes, *Gpx3*, showed expression restricted to the four CO GE clusters but no expression in any of the DES clusters, providing further evidence that DES exposed mice lack normal GE. These findings provide a panel of markers that can be used for the identification of uterine GE cells at all stages of differentiation.

### Disruption of uterine epithelial stem/progenitor cells in DES-exposed uteri

To identify epithelial stem cells (EpSC) in CO epithelium clusters, we used markers previously reported as expressed in uterine stem cells (*Aldh1a1, Lgr5, Axin2*)^16,38–40^. Dual feature plots revealed high expression of all three markers in CO cluster 12, with some cells expressing combinations of these genes (Figure 5A). *Aldh1a1* and *Axin2* were also expressed in mature GE, while *Lgr5* was also expressed in developing GE. There was limited expression of *Axin2* in the developing GE. Because the uterine EpSC population has a very slow turnover rate^17^, we used feature plots to identify clusters with high expression of cell division markers *Top2a* and *Mki67*. Only a few cells in the putative EpSC population had any *Top2a* or *Mki67* expression (Figure 5A). However, both markers were highly expressed in the GPC (CO cluster 14) and in CO cluster 4, which almost entirely lacked *Aldh1a1* expression (Figure 5A; Table S5A). We interpret these results to indicate that CO cluster 4 consists of luminal progenitor cells (LPC) and CO cluster 12 consists of EpSC. An overlap of the 596 GPC up-regulated DEGs with the 502 LPC up-regulated DEGs showed 225 in common, suggesting they are related but distinctly different cell populations (Tables S8A-C).

To determine if the DES epithelial clusters exhibited normal EpSC, GPC or LPC expression patterns in specific clusters, dual feature plots of *Aldh1a1, Lgr5, Axin2, Top2a* and *Mki67* were generated (Figure 5A). *Aldh1a1* was expressed in rare cells in DES cluster 4; however, it was not differentially expressed in any DES cluster (Table 5B). In addition, *Aldh1a1* expression was not overlapped by *Lgr5* or *Top2a*, suggesting these *Aldh1a1+* cells were not EpSC or progenitor cells. *Lgr5* and *Axin2* were present in most DES epithelial cell clusters with neither being DES cluster DEGs (Figure 5A; Table S5B). Both cell proliferation markers *Top2a* and *Mki67* were highly expressed in DES cluster 6, with *Top2a* also having sporadic expression in DES cluster 4, for which it was identified as a DEG (Figure 5A; Table S5B). These data suggest that there was only one progenitor cell population in the DES epithelial clusters, predominantly comprised of DES cluster 6. However, we did not clearly identify any EpSC in the DES cells.

Because we failed to identify EpSC in the DES clusters, we returned to the integrated (CO + DES) epithelial cell analysis (Figure 3A) to identify the DES clusters mapping most closely to CO EpSC. The CO EpSC in the integrated UMAP were identified by their expression of *Aldh1a1* in an integrated feature plot, with strong expression in only two regions: in a few GE cells and in a small set of cells adjacent to the large non-basal cell DES clusters (Figure 5B). The CO EpSC were located within integrated clusters 7 and 18, which were mainly comprised of DES cells, but most DES cells in these clusters expressed *Aldh1a1* at a low level or not at all (Figure 5B). The UMAP locations of the large set of non-basal DES epithelial cells near CO EpSC and far from the CO LE and CO GE indicates that even though DES EpSCs could not be identified, the non-basal DES epithelial cells were more closely related to CO EpSC than to LE or GE cells.

To test whether our identification of the stem and progenitor cell clusters was consistent with their spatial localization, we localized ALDH1A1 and TOP2A in uterine tissue. In CO uteri, ALDH1A1 was highly expressed in glands farthest away from the LE, consistent with its expression in mature GE and confirmed by the location of FOXA2 immunostaining in adjacent sections (Figures 5C and S4)^16,39^. ALDH1A1 was also expressed sporadically throughout the CO LE, consistent with previous reports of the epithelial stem cells residing in this location^17^. In DES uteri, ALDH1A1 was occasionally expressed in the deepest GE, similar to controls; however, the vast majority of GE had no detectable ALDH1A1 (Figure 5C). In contrast to CO, there was no ALDH1A1 detected in the LE of the DES exposed uteri. These data suggest that the *Aldh1a1* expression observed in non-integrated DES cluster 4 was from the GE cells. In CO, TOP2A was expressed sporadically across the LE as well as in invaginating GE, the region previously reported to contain epithelial stem/progenitor cells, supporting the scRNAseq identification of two distinct populations of CO progenitor cells, GPC and LPC (Figure 5D)^17,18^. In agreement with the scRNAseq cluster analysis, ALDH1A1 and TOP2A protein expression was completely non-overlapping (Figure 5C). In DES uteri, TOP2A was expressed in many LE and GE cells. The presence of *Top2a+/Mki67+* cells only in DES cluster 6 suggests that these cells lack GE or LE identity, despite their localization in both epithelial regions in uterine tissue. ALDH1A1 and TOP2A immunostaining in serial sections of DES-exposed uteri clearly demonstrated that these two proteins did not overlap (Figure 5C), confirming the lack of *Aldh1a1* in the DES progenitor cell population.

In addition to these known uterine stem cell and progenitor cell markers, two of the highest DEGs in the cluster of CO EpSC encode proteins in the Wnt/β-catenin signaling pathway, engrailed 2 (*En2*) and *Wnt7a* (Tables 5A and 6C). *Wnt7a* is a secreted signaling molecule that regulates female reproductive tract differentiation including GE development, serving as a cell death suppressor in the uterus^41,42^. *En2* is a homeobox transcription factor that can repress *Wnt* signaling but is also an oncogene^43,44^. The top DEG in CO EpSC was *Gstm7*, and three additional *Gstm* genes were DEGs in this cluster (Table S6C). These genes encode mu family glutathione-S-transferases, which function to detoxify electrophilic molecules including products of oxidative stress^45,46^. Dual feature plots of *En2*, *Wnt7a*, and *Gstm7* in CO showed high expression of all three in EpSCs and sporadic expression in LPC and GPC clusters (Figure 5D). *En2* was also highly expressed in a few cells of CO LE cluster 5. In contrast, all three genes were expressed in most DES clusters including DES cluster 6, the progenitor cell population, though they mainly were not expressed in the same cells (Figure 5D). In CO uteri, EN2 protein was quite restricted, with highest expression at the edge of budding GE and some expression in LE near these regions (Figure 5D). There was very low expression of EN2 in the CO GE. In DES uteri, EN2 was expressed in most epithelial cells including LE and GE, with some GE expressing very high levels. These data demonstrated a restricted pattern of stem/progenitor cell gene expression in normal uterine epithelial cells and the loss of this restriction in DES uteri, resulting in most epithelial cells expressing several stem/progenitor cell-specific genes.

The excessive aberrant expression of *Wnt7a* and Wnt signaling targets *Axin2* and *En2* in the DES cells suggested additional members of this pathway could be similarly disrupted. Violin plots of select Wnt signaling pathway ligands, receptors and targets demonstrated moderate or high expression of most these genes in the CO EpSC population (Figure 5E). Other CO cell types had very restricted expression patterns. For example, *Wnt4, Fzd1* and *Numb* were restricted to LE cell populations and *Wnt7b, Fzd10, Myc* and *Hdac11* were restricted to developing GE. These data demonstrated tight regulation of Wnt ligands/receptors and their downstream targets in a cell type specific manner. In DES clusters, there was widespread expression of most Wnt ligands, receptors, and targets (Figure 5E). These data show that DES epithelial cells exhibit severe disruption of the normally restricted expression pattern of Wnt signaling pathway genes and suggest widespread activation of Wnt signaling.

### DES induced uterine cancer is characterized by activation of Wnt and PI3K/AKT signaling pathways

To identify cancer cells in the non-basal cell DES clusters, we used expression of *Six1*, a reliable biomarker of neonatal DES exposure-induced cancer^14,20,26,27^. *Six1* was most highly expressed in DES cluster 5 (Table S5B). The cancer-associated genes *Olfm4* and *Rad51b* were among the top DEGs in this cluster. *Olfm4* is both an oncogene and a target of Wnt signaling that provides negative feedback to this pathway, and *Rad51b* is a marker of DNA damage^47–49^. There was minimal sporadic expression of these markers in CO epithelial cells (Figure 6A). DES cluster 5 epithelial cells had high overlapping expression of *Six1* and *Olfm4* and almost exclusive expression and extensive overlap of *Olfm4* and *Rad51b* (Figure 6A). A feature plot of *Olfm4* in the spatial transcriptomics uterine tissue sections showed high expression in cancer regions in the DES sample and a lack of expression in the CO (Figure 6B).

**Figure 6:**
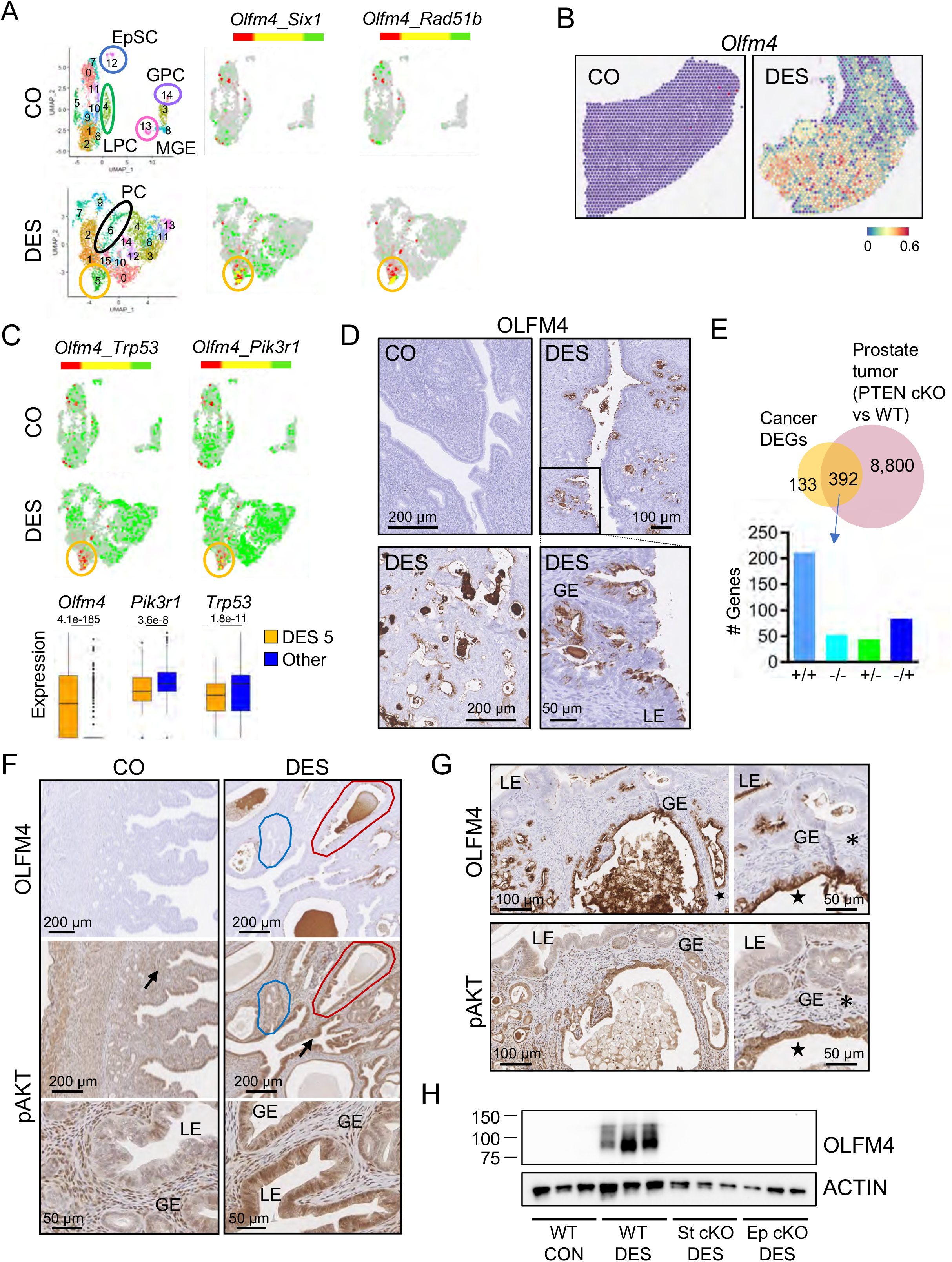
Cancer cells in DES uteri are characterized by OLFM4 expression and PI3K/AKT signaling. A) UMAPs of non-integrated CON and DES epithelial cells. EpSC (blue), GPC (purple), LPC (green) and MGE (pink) are indicated in CO (top). PC (black) and cancer population (gold) are indicated in DES (bottom). Dual feature plots of cancer cell markers (*Olfm4, Six1* and *Rad51b*) for CO (top) and DES (bottom). B) Expression of *Olfm4* in spatial transcriptomics sections; CO (left), DES (right). Expression levels are natural log transformed counts. C) Dual feature plots of select cancer cell markers in CO (top) and DES (bottom). Cancer population, gold circle. Box and whisker plots of cancer markers for DES cluster 5 versus all other DES clusters. Expression levels are natural log transformed counts; adj p-values from Table S5B are indicated for each gene. D) OLFM4 immunohistochemistry in CO and DES sections, as indicated. Bottom right section is magnified view of boxed region in top right section. Panels D-F: LE, luminal epithelium; GE, glandular epithelium. E) Venn diagram of DES cluster 5 DEGs (cancer population) and prostate tumor DEGs [prostate PTEN conditional KO (cKO) versus wild type (WT)]. Data generated from BaseSpace Correlation Engine knockout atlas (Illumina); original data from ^54^. Overlapping genes categorized as increased (+) or decreased (-) in each study and split into four categories (+/+, −/−, +/− or −/+; DES cluster 5 DEGs listed first, prostate tumors listed second) on graph. F) OLFM4 and pAKT immunohistochemistry in adjacent uterine tissue sections from 12-month-old CO (left) and DES (right). Blue outline, region of low OLFM4/pAKT; red outline, high OLFM4/pAKT. Arrows indicate pAKT stained regions shown at higher magnification below. G) OLFM4 (top) and pAKT (bottom) immunohistochemistry in adjacent uterine tissue sections from 9-month-old FVBN/J DES exposed mice. Higher magnifications from same sections (right). Asterisks, OLFM4-/pAKT-cells; stars, OLFM4+/pAKT+ cells. H) Immunoblot of OLFM4 in uteri from 12-month-old control or DES-treated mice. Each lane has 5 μg uterine protein extract from one mouse (n=3 mice per group). Actin used as a loading control. WT, wild type; St cKO, stromal ERα conditional knockout; Ep cKO, epithelial ERα conditional knockout.

Activating mutations in β-catenin, which are accompanied by activation of Wnt signaling, are associated with human endometrioid endometrial carcinomas^50^. This observation, combined with our previous data showing widespread activation of Wnt signaling in DES epithelial cells, suggested that Wnt signaling was driving the DES cancer phenotype. Examination of DES cluster 5 for expression of Wnt signaling targets, however, indicated that relative to other DES clusters, the DES cancer cells had among the lowest expression of Wnt ligands, receptors, and target genes including *Wnt7a, Ccnd2, Axin2*, and *Myc* (Figure 5F). These findings were inconsistent with Wnt signaling as the sole cancer driver.

In the mouse, loss of the tumor suppressor PTEN in the uterus leads to rapid development of endometrial cancer, even if it is only deleted in the epithelial cells, and combined loss of PTEN and a second tumor suppressor, TRP53, induces higher grade cancer with invasion^51,52^. *Trp53* and *Pik3r1*, which encodes a PTEN-stabilizing protein^53^, were widely expressed in most CO and DES clusters but were downregulated in DES cluster 5 (Figure 6C; Table S5B). Dual feature plots showed an inverse relationship between *Olfm4* and both *Trp53* and *Pik3r1;* the differences between expression of these 3 genes in DES cluster 5 relative to all other clusters were highly significant (Figure 6C). To confirm the identification of DES cluster 5 as the cancer cell population, we performed OLFM4 immunohistochemistry. OLFM4 was not detected in CO uterine tissue but was robust in cancer regions in DES uterine tissue (Figure 6D). This expression was variable across animals, with a direct correlation between extent of uterine cancer and extent of OLFM4 staining (Figures 6D and S5A). Of note, OLFM4 staining was present in both LE and GE (Figures 6D and S5A). These data confirmed the identity of the uterine cancer cell population in the scRNAseq dataset and identified *Olfm4* and *Rad51b* as additional markers of this cancer type.

The reductions in expression of *Trp53* and *Pik3r1* in the cancer cell population suggested a role in DES cancer formation for increased PI3K/AKT signaling due to loss of PTEN. To further examine the molecular signature of the DES cancer cells, we compared the DES cluster 5 DEGs to curated gene perturbation models in the BaseSpace Correlation Engine knockout atlas (Illumina). The gene perturbation with the highest correlation to this dataset was PTEN. One dataset with high overlap was a conditional deletion of *Pten* in prostate epithelium, which results in prostate tumors^54^ (Figure 6E; Table S9). Of the DES cluster 5 DEGs, 392/525 (75%) overlapped the DEGs in this prostate cancer model; 67% of these genes were altered in the same direction (Figure 6E). These data suggest that loss of PTEN activity plays a major role in the formation of neonatal DES exposure-induced uterine cancer.

To test whether PI3K/AKT signaling was activated in DES induced uterine cancer cells, we localized OLFM4 and phosphorylated AKT (pAKT) in adjacent sections. CO uteri (during estrus) had diffuse pAKT immunoreactivity in most cell types but only sporadically in the nuclei of LE cells and lower staining in GE compared to stroma (Figure 6F). DES uteri had higher pAKT nuclear and cytoplasmic immunoreactivity in LE cells compared to CO and there was very strong pAKT nuclear and cytoplasmic staining in most GE. The regions of more intense pAKT staining generally corresponded with regions of OLFM4 expression (Figures 6F and S5). Because DES-exposed CD-1 mice only develop localized foci of uterine adenocarcinoma by 12 months of age, we further explored the correlation of OLFM4 and pAKT in a more robust model of developmental DES-induced uterine cancer. Female FVBN/J mice develop uterine adenocarcinoma at an earlier age (6 months) and they exhibit very extensive adenocarcinoma throughout the uterus by 9-12 months of age following the same neonatal DES exposure paradigm used in our standard CD-1 mouse model^27^. Uterine tissue sections from 9-month-old DES exposed FVBN/J mice were immunostained for OLFM4 and pAKT in adjacent sections. As previously observed, FVBN/J mice exhibited more extensive cancer regions. These regions had more robust OLFM4 expression relative to that in CD-1 mice (Figure 6F, 6G, S5A and S5B). Staining of adjacent sections for pAKT revealed consistent overlap of pAKT with OLFM4+ cancer regions (Figure 6G).

The data presented so far suggests a model in which abnormally differentiated epithelial cells are influenced by stromal inflammation and oxidative stress to activate PI3K/AKT signaling that then drives uterine adenocarcinoma development. However, we have not yet demonstrated that DES-induced changes in epithelial or stromal gene expression are essential for the cancer phenotype. To answer this question, we generated conditional knockout (cKO) mice lacking ERα only in endometrial stromal cells or only in endometrial epithelial cells^55,56^. The loss of ERα in these tissue compartments precludes DES action on the targeted cell type. Uteri were collected from 12-month-old control mice and neonatal DES exposed wild type, stromal ERα cKO, and epithelial ERα cKO mice and the presence of uterine cancer tested by immunoblotting for OLFM4. DES exposed wild type mice had high levels of OLFM4, but neither controls nor either of the DES-exposed cKO lines expressed this cancer marker (Figure 6H). These findings indicate that DES-induced changes in cellular differentiation pathways of both epithelial and stromal cells are required for the development of uterine adenocarcinoma.

## Discussion

The dataset reported here provides a rich resource of scRNAseq information from over 48000 total adult mouse uterine cells, allowing an in-depth analysis of cell types. Previous scRNAseq analyses of rodent uterine cells reveal interesting information regarding postnatal uterine development^16,25,28^ and pregnancy^57,58^. Two datasets contain cells from adult non-pregnant uterus, but both are limited by having few cells for analysis^16,59^. Here we focused mainly on epithelial cells because they are the cell type that develops into uterine adenocarcinoma following a developmental insult. The large number of epithelial cells from adult control females in estrus enabled clear identification of stem cells, progenitor cells, luminal cells, and glandular cells, including a glandular cell developmental trajectory.

Our findings confirm previous observations that uterine EpSC are localized at the intersection between glands and lumen^17^. However, instead of EpSCs differentiating into a single progenitor cell population^17^, we show that they differentiate into two distinct proliferating progenitor cell types, one destined to become luminal epithelium and one to become glandular epithelium (Figure 7). The EpSCs are marked by high expression of EN2 protein and *Wnt7a*, but neither marker is entirely specific to stem cells, and the presence of ALDH1A1 in mature glands argues against its previously suggested specificity as a marker of stem/progenitor cells^16^. Excitingly, we identified a new uterine EpSC marker, *Gstm7*, and found that several additional *Gstm* isoforms were highly expressed in EpSC. This finding mirrors previous observations of increased *GSTM* family gene expression in human fetal liver hematopoietic stem cells, including *GSTM2*, the mouse *Gstm7* homolog^60^. Enrichment of enzymes that function in a pathway responsible for managing and eliminating reactive oxygen species from the cell makes sense for uterine EpSCs, which must be preserved for the lifetime of the individual.

**Figure 7:**
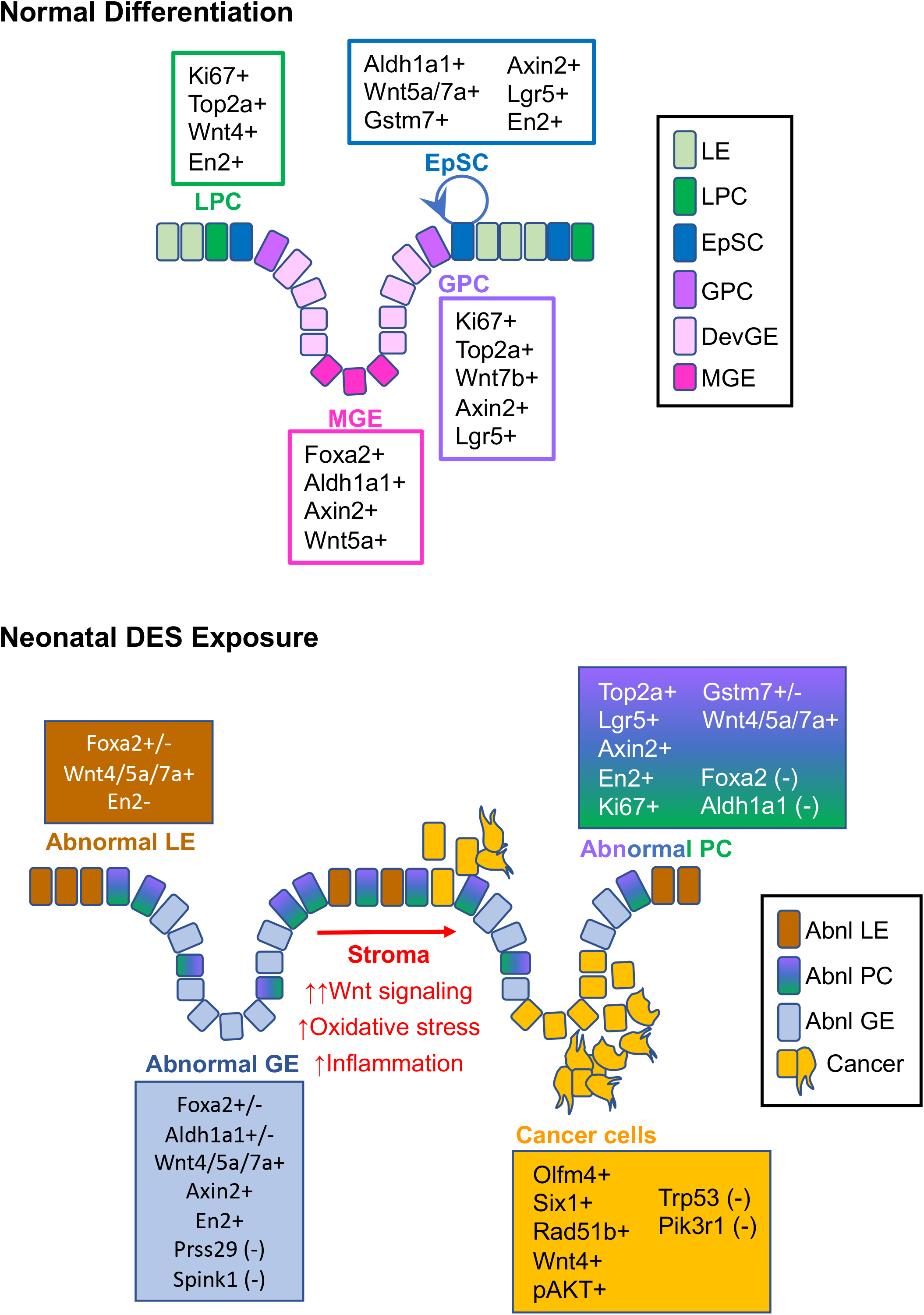
Model of uterine epithelial cell differentiation in control and DES-exposed uteri. Normal differentiation: Rare epithelial stem cells (EpSC) transition to proliferative luminal progenitor cells (LPC) or glandular progenitor cells (GPC). LPC transition to luminal epithelium (LE) and GPC transition to developing glandular epithelium (DevGE) and then mature glandular epithelium (MGE). Neonatal DES exposure: Proliferative abnormal progenitor cells (Abnl PC) transition to either abnormal luminal epithelium (Abnl LE) or abnormal glandular epithelium (Abnl GE), which are distinguished mainly by their histological locations. Widespread activation of Wnt signaling in the presence of stromal oxidative stress and inflammation activate PI3K/AKT signaling to drive malignant transformation of abnormal epithelial cells. Markers present at the various stages of differentiation for control and DES-exposed uteri are indicated.

DES exposure permanently disrupts differentiation of uterine epithelium. The abnormal epithelial cells have characteristics of luminal and glandular epithelium as well as EpSCs, but no distinct EpSC population. Instead, we found a single progenitor cell population that could not be further characterized as glandular or luminal, suggesting substantial problems with the downstream fate of these cells. The abnormal epithelial differentiation trajectory was confirmed by the constitutive activation of Wnt signaling in most DES epithelial cells, in stark contrast to the epithelial subtype specificity observed in differentiated CO epithelial cells. Increased Wnt signaling due to activating mutations in pathway members can cause human endometrial cancer^61,62^. However, it is unlikely that Wnt signaling alone explains neonatal DES-induced uterine cancer because persistent activation of Wnt signaling in the mouse uterus using genetic approaches results in uterine hyperplasia but not cancer^63,64^. In addition, the DES cancer cell cluster had reduced Wnt pathway activation relative to the other DES epithelial cell clusters. Instead, this cluster had diminished *Trp53* and *Pik3r1* mRNAs and elevated levels of PI3K/AKT signaling, which are common drivers of many cancer types, including endometrial cancer in humans and animal models^51,61,65–67^. The impact of abnormal stromal cell inflammatory and oxidative stress responses on the adjacent epithelium cannot be ignored in this model because both DES initiated stromal and epithelial changes are required for cancer development. Overall, these findings provide compelling evidence that abnormal activation of PI3K/AKT signaling is the long-sought explanation for uterine cancer development following developmental estrogen exposure (Figure 7).

The DES model has a striking resemblance to a mouse model of intestinal tumorigenesis induced by expression of a constitutively active form of β-catenin in intestinal epithelial cells (IEC)^68^. In the IEC model, active β-catenin induces dedifferentiation of epithelial cells, which gain stem cell characteristics. β-catenin induces activation of the inflammatory mediator nuclear factor-κB (NF-κB), which serves to amplify β-catenin-mediated signaling and induction of IEC tumor formation. Similarly, the DES model has incomplete differentiation of epithelial cells associated with widespread activation of Wnt signaling, combined with inflammation in the stroma as a cell proliferation signal. The main difference between the two models is that the IEC tumor model requires genetic manipulation to drive β-catenin signaling, whereas the DES model simply requires estrogen-mediated signaling for a brief period during epithelial cell differentiation. The similarity between these models highlights the influence of environmental exposures during development on tumor susceptibility later in life.

An important question remains: how does PI3K/AKT signaling become activated in the cancer cells? Because most of the epithelial cells, not just the cancer cells, have persistent activation of Wnt signaling, it is likely that this signaling pathway is activated because of the “first hit” of DES exposure and sets the stage for later activation of PI3K/AKT in the cells that become malignant. There are many connections between Wnt and PI3K/AKT signaling pathways, including common components relevant to cancer development such as MYC, GSK3, PTEN, and CCND1 [reviewed in ^64^]. Recently, WNT3A-mediated activation of PI3K was demonstrated in colorectal cancer cells^69^. Additional connections between these pathways could be mediated by noncoding RNAs such as microRNAs or long noncoding RNAs^70^. Finally, stromal inflammation- and oxidative stress-induced genotoxicity and cell proliferation may promote cancer development through activation of NF-κB and PI3K/AKT signaling in epithelial cells^71,72^. NF-κB appears to be activated in the DES cancer model based on pathway analysis of altered genes, including *Olfm4*, which can be upregulated by NF-κB, among other transcription factors^48,73^. A bidirectional interaction between the inflammatory stroma and the Wnt-activated epithelial cells likely contributes to cancer development in this model.

The most frequently mutated genes in human endometrial endometrioid cancer include the tumor suppressors TP53, PTEN, and PIK3R1; the overwhelming majority of these cancers have abnormal activation of the PI3K/AKT signaling pathway^53,61,66^. In human endometrial organoids, inhibition of Wnt signaling increases epithelial cell differentiation^74^, suggesting that persistent Wnt signaling in human endometrium, as in the mouse, could reduce epithelial differentiation. Instead of a primary mutational event, perhaps some human endometrial cancers result from an early environmental insult during cellular differentiation, leading to Wnt pathway activation and abnormal differentiation of epithelial cells. Later induction of PI3K/AKT signaling could be related to stromal inflammation and subsequent induction of mutational events through increased oxidative stress, downregulation of *Pi3kr1* mRNA, or other alterations in regulators of the PI3K/AKT pathway.

## Acknowledgements

We thank Ciro Amato, Humphrey Yao, and Guang Hu for critical review of the manuscript. We are grateful to Franco DeMayo for providing the Amhr2-cre mouse line and Ken Korach for providing the Wnt7a-cre and Esr1-floxed mouse lines. This work was supported by the Intramural Research Program of the National Institutes of Health, National Institute of Environmental Health Sciences (1ZIAES102405).

## Author contributions

E.P.-B., W.N.J., A.A.S. and C.J.Williams designed the study. E.P.-B., W.N.J., A.A.S., X.X., and D.V.C. conducted the experiments. B.N.P. performed the bioinformatic analyses. C.J.Willson and E.N.Q. performed histological analysis of spatial transcriptomics samples. E.P.-B., W.N.J., B.N.P. and C.J.Williams wrote the first draft of the manuscript. All authors edited the manuscript.

## STAR ✶ METHODS

### RESOURCE AVAILABILITY

#### Lead Contact

Further information and requests for resources and reagents should be directed to and will be fulfilled by the Lead Contact, Carmen Williams.

#### Materials Availability

This study did not generate new unique reagents.

#### Data and Code Availability

The sequencing data generated in this study have been deposited in the Gene Expression Omnibus database under accession code GSE218156 and will be publicly available as of the date of publication. This paper does not report original code. Any additional information required to reanalyze the data reported in this paper is available from the Lead Contact upon request.

### EXPERIMENTAL MODEL AND SUBJECT DETAILS

Animal care guidelines under an approved protocol for the National Institutes of Health were strictly followed. CD-1 mice were obtained from the NIEHS in-house breeding colony and housed under conditions previously reported^29^. FVBN/J mice were purchased from The Jackson Laboratory (Jax#001800) to generate female pups for experiments described below. Previous work demonstrated that C57BL/6 mice do not develop uterine adenocarcinoma; therefore, we first generated mouse lines for ERα (*Esr1*) flox, Amhr2-cre and Wnt7a-cre that were backcrossed to an FVBN/J background for ten generations^55,56,75^. ERα (*Esr1*) conditional knock out (cKO) mice were generated by crossing Esr1 floxed mice (FVBN/J background) with either Amhr2-cre mice (stromal cKO; FVBN/J background) or Wnt7a-cre mice (epithelial cKO; FVBN/J background); proper names are listed in the Key Resources2Table.

### METHOD DETAILS

#### Animal treatments and tissue collection

Briefly, female pups delivered from timed pregnant dams listed above were randomly distributed to 10 female pups per litter and then randomly assigned treatment groups of either corn oil (CO) or DES (1 mg/kg/day). Pups were exposed on neonatal days 1-5 by subcutaneous injection using a volume of 0.02 mL; mice were weaned and housed as described previously^29^. CD-1 mice exposed to DES at this dose and timing develop uterine adenocarcinoma (incidence >50%) at 12 months of age so we collected uteri for single cell isolation and frozen tissue sections at this age^14^. Adult DES exposed mice are in persistent estrus; therefore, CO mice in estrus were selected using vaginal observation^76,77^. Uteri from FVBN/J mice were collected at 9 months of age for immunohistochemistry. For immunoblotting, uteri were collected at 12 months of age from control and DES-exposed wild type (Esr1 flox/flox), epithelial cKO (Esr1 flox/flox, Wnt7a-cre+) and stromal cKO (Esr1 flox/flox, Amhr2-cre+) mice and frozen at −80°C until use.

#### Single cell isolation from adult uterine tissue

Mice were euthanized and uterine horns were excised away from the uterine body and the oviducts. Uteri from two CO and four DES mice were pooled for single cell isolation. All procedures were performed on ice unless otherwise specified. Uteri were rinsed in calcium and magnesium free phosphate buffered saline (PBS-CMF), horns were slit open lengthwise and soaked in PBS to remove any debris. Cell dissociation was performed by incubating tissue in Trypsin-EDTA (0.25%) (Gibco-Thermo Fisher) on ice for 1 hour, then 10 min at room temperature, then 50 min in a 37°C water bath with gentle agitation. Tissue was removed from media, and the cell suspension was centrifuged at 450 x *g* for 10 min at 4°C. Pellets were resuspended in DMEM/F12 (Gibco-Thermo Fisher) containing 10% heat inactivated fetal bovine serum (FBS; Thermo Fisher), serially filtered through sterile CellTrics 100 μm and 30 μm filters (Fisher Scientific) and centrifuged at 450 x *g* for 10 min at 4°C. Pellets were resuspended in 1-2 mL PBS-CMF containing 0.04% AlbuMAX bovine serum albumin (BSA; Thermo Fisher). A small aliquot of cell suspension was stained with Trypan Blue (Gibco-Thermo Fisher) and counted using a hemocytometer.

#### Single cell RNA sequencing

Single-cell libraries were prepared using the Chromium platform with the Chromium Single Cell 3’ Reagent Kit v3 (Cat. 1000075, 10x Genomics Inc.) following the manufacturer’s protocol. Briefly, freshly prepared single cells and single gel beads conjugated with cell barcodes and reverse transcription primers were partitioned into oil droplets as emulsion in the 10x Genomics Chromium Controller instrument followed by cell lysis and barcoded reverse transcription of mRNA, cDNA amplification by PCR, fragmentation, and adding adapters and sample index amplification by PCR. Libraries were sequenced on an Illumina NovaSeq 6000 for paired end reads: read 1, 30 bp; read 2, 100 bp.

The scRNA count matrix was generated using cellranger v3.0.1, using GENCODE genes for mm10 (downloaded from 10X Genomics Inc.’s website on 3/22/2018), which were filtered according to 10x Genomics recommendations. Potential barcode swapping was identified using the R package DropletUtils v1.14.2, and potential cell doublets were identified and removed using the R package scran v1.22.2. Downstream analysis was carried out using the R package Seurat v3.1.0 following the standard SCTransform-based pipeline with MAST used for differential expression testing.

#### Spatial Transcriptomics

The Visium Spatial Gene Expression Slide kit (10x Genomics, Cat. 1000184) was used for the Spatial Transcriptomics study. Uterine tissue freezing and embedding was performed following the 10x Genomics Visual Spatial Protocol-Tissue Preparation Guide, CG000240-Rev A. Briefly, uterine tissues were frozen in isopentane and embedded in chilled Tissue-Plus O.C.T. compound (Fisher Scientific) on dry ice. Tissue sections were stained with Mayer’s hematoxylin (Millipore Sigma) and eosin (Millipore Sigma) (H&E) and evaluated by a board-certified veterinary pathologist for the presence of luminal and glandular epithelial cells in both CO and DES sections and the presence of uterine adenocarcinoma in the DES sections (n=5 mice per group). Two mice per group were selected for subsequent sectioning for spatial transcriptomics. Selected blocks were cored to capture the areas of interest and fit the capture area on the Gene Expression slide. Tissues were cryo-sectioned at 10 μm thickness and immediately placed on the Gene Expression slide. After four areas were captured (two control and two DES) the slide was kept at −80°C until staining. The slide was fixed with methanol at −20°C for 30 min and stained with H&E per manufacturer’s instructions. The slide was imaged using an Aperio AT2 slide scanner (Leica Biosystems). Optimal permeabilization time of 18 min was determined using the tissue optimization kit (10x Genomics, Cat. 1000193). On-slide mRNA reverse transcription, cDNA synthesis and release, cDNA amplification, and library preparation were performed following instructions in the user manual. The libraries were then sequenced on a NextSeq 500 for paired end reads: read 1,28bp; read 2, 90bp. The spatial sequencing data was processed using 10X Genomics Space Ranger v1.2.2 and analyzed using Seurat v4.0.5. The reference genome used for the scRNAseq analysis was mm10.

#### Immunohistochemistry

Whole uterine tissues from CO and DES mice (n=4-6 mice per group) were fixed in 10% neutral buffered formalin for ~48 hours, changed into 70% ethanol, processed and embedded in paraffin blocks and sectioned at 6 μm thickness. Briefly, slides were deparaffinized in xylene, rehydrated through graded ethanol, and blocked for endogenous peroxidases using 3% hydrogen peroxide for 15 min. Immunohistochemistry methods have been described in detail previously^78^. All concentrations and detailed protocol instructions for each antibody can be found in Table S10. Protein/antibody complexes were visualized by using 3-diaminobenzidine (DAB) chromagen (Dako) for six minutes, counterstained with hematoxylin, dehydrated and cover slipped. Slides were scanned using an Aperio AT2 slide scanner (Leica Biosystems). Images were captured using Aperio ImageScope v. 12.4.3.5008 (Leica Biosystems).

#### Immunoblotting

Uterine tissues were pulverized on dry ice and total protein isolated using TPER (Thermo Fisher). Protein concentration was assessed using a Qubit protein assay (Thermo-Fisher). Samples (5 μg) were loaded on a 10% TGX gel (Bio-Rad), run at 150 V and transferred to PVDF membrane using the Trans-Blot Turbo transfer system (Bio-Rad). The blot was blocked in 5% Blotto (Thermo Fisher) in tris buffered saline plus 0.1% Tween-20 (TBS-T) for 1 h at RT. Rabbit monoclonal anti-OLFM4 (Cell Signaling Technology) was diluted to 0.6 μg/mL in 5% blocking solution and applied to the blot overnight at 4°C. The blot was washed three times 15 minutes with TSB-T, incubated with donkey anti-rabbit IgG diluted 1:25,000 in 1% blocking buffer for 1 h at RT. Following three washes in TBS-T for 15 min each, immunoreactive bands were visualized using Super Signal West Femto reagents (Thermo-Fisher) following the manufacturer’s instructions. Images were captured using a ChemiDoc Touch Gel Doc system (Bio-Rad). Antibodies were stripped with Restore (Thermo-Fisher) for 30 min at 37°C.

Peroxidase conjugated mouse monoclonal anti-β-actin (Sigma) at a dilution of 1:10000 in 5% blocking solution was applied for 1 h at RT and visualized as described above.

### QUANTIFICATION AND STATISTICAL ANALYSIS

For immunohistochemical analysis, a minimum of 4-6 mice per group were immunostained with each antibody and representative images are shown. For immunoblotting, three mice per group were tested for OLFM4 – all samples are shown in immunoblot in Figure 6H.

For scRNA-seq, statistical analysis was carried out using the R package Seurat v3.1.0 following the standard SCTransform-based pipeline with MAST used for differential expression testing. For spatial transcriptomics, statistical analysis was carried out using 10X Genomics Space Ranger v1.2.2 and analyzed using Seurat v4.0.5. p-values are included in the supplementary tables generated for each analysis presented. Statistical analysis for data presented as box and whisker plots in Figure 6C are from Table S5B.

## Supplemental tables not included in main supplemental PDF

Table S1A: Raw data for single cells captured by single cell RNA-seq

Table S1B: Differentially expressed genes (DEGs) that are specific to each cell type

Table S2A: DEGs of CO spatial transcriptomics clusters

Table S2B: DEGs of DES spatial transcriptomics clusters

Table S3A: Stromal genes in CO cluster 8 not found in DES cluster 4

Table S3B: Stromal genes in DES cluster 4 not found in CO cluster 8

Table S3C: Stromal genes in common between CO cluster 8 and DES cluster 4

Table S3D: GO categories for 302 CO stromal specific genes (p<0.01, gene count≥10)

Table S3E: GO categories for 339 DES stromal specific genes (p<0.01, gene count≥10)

Table S3F: Non-overlapping GO categories for CO stromal specific genes

Table S3G: Non-overlapping GO categories for DES stromal specific genes

Table S4: Integrated CO and DES epithelial cell clusters

Table S5A: DEGs of CO non-integrated epithelial cell clusters

Table S5B: DEGs of DES non-integrated epithelial cell clusters

Table S6A: GE specific DEGs

Table S6B: LE specific DEGs

Table S6C: CO cluster 12 specific DEGs

Table S7A: DEGs from Slingshot analysis of CO GE

Table S7B: GE only DEGs from Slingshot analysis

Table S8A: GPC CO cluster 14 specific DEGs

Table S8B: LPC CO cluster 4 specific DEGs

Table S8C: 225 DEGs in common between LPC and GPC

Table S9: Uterine cancer (DES cluster 5 DEGs) overlapped with Pten conditional knock out prostate tumors versus normal prostate

Table S10: Antibodies used in the current study

## KEY RESOURCES

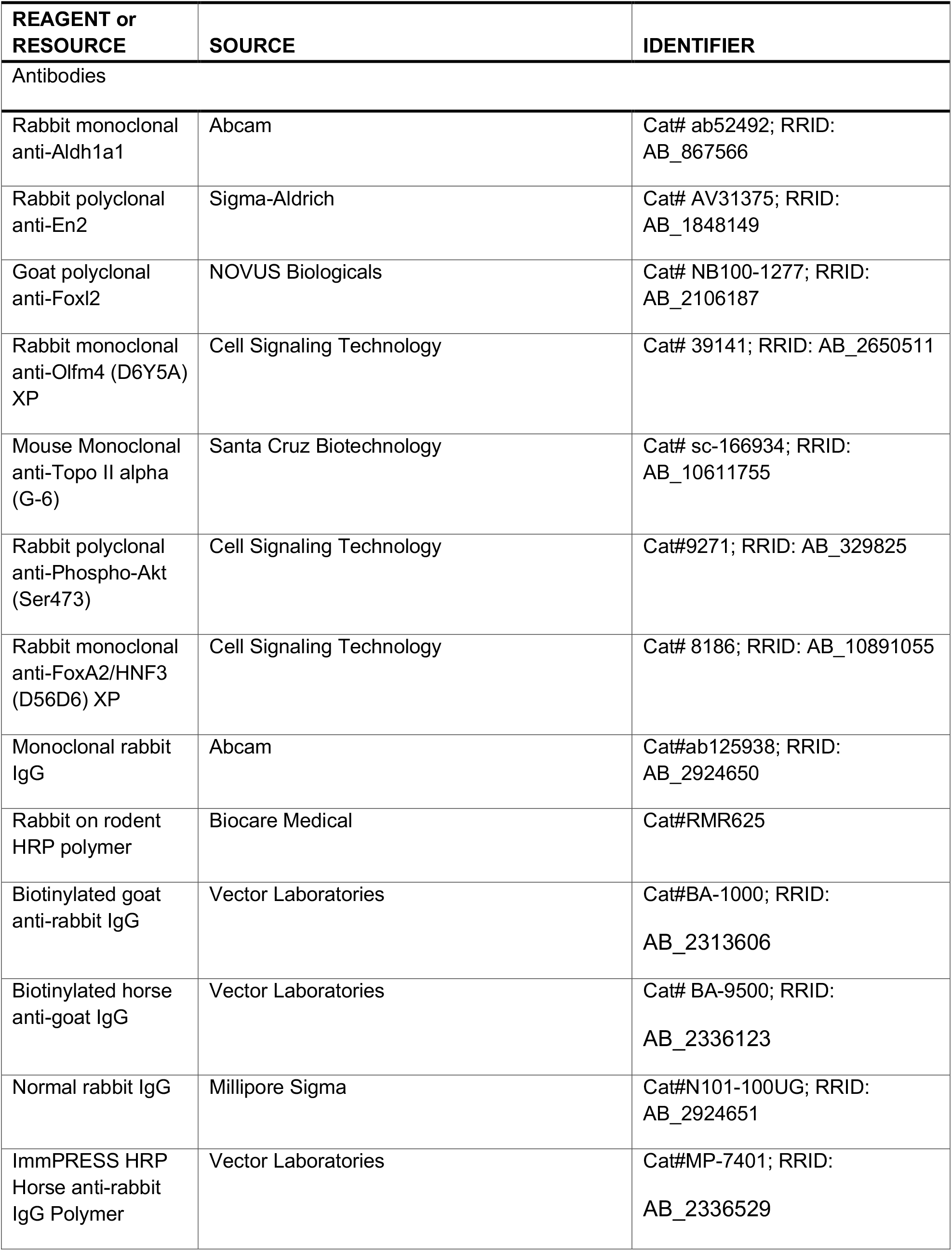
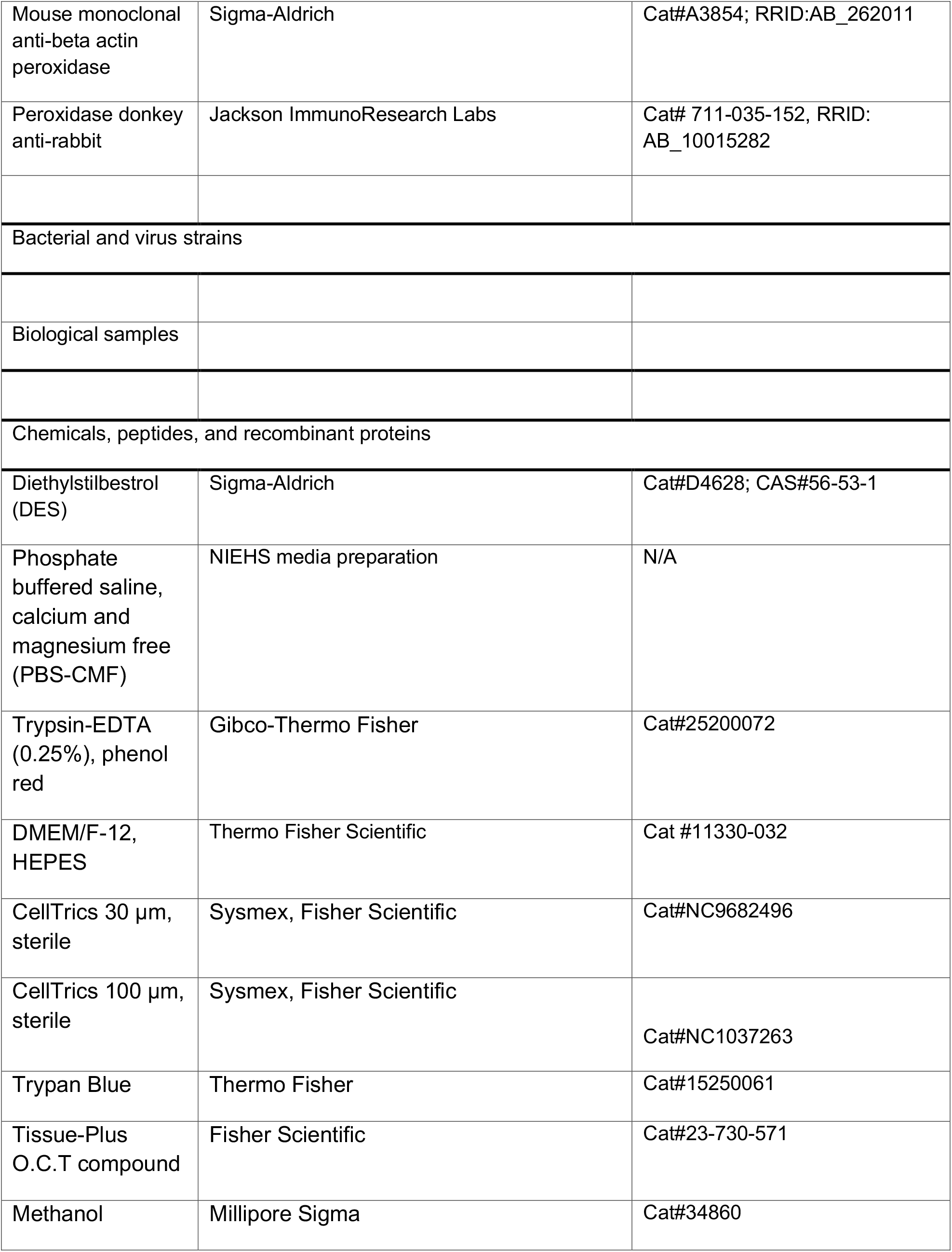
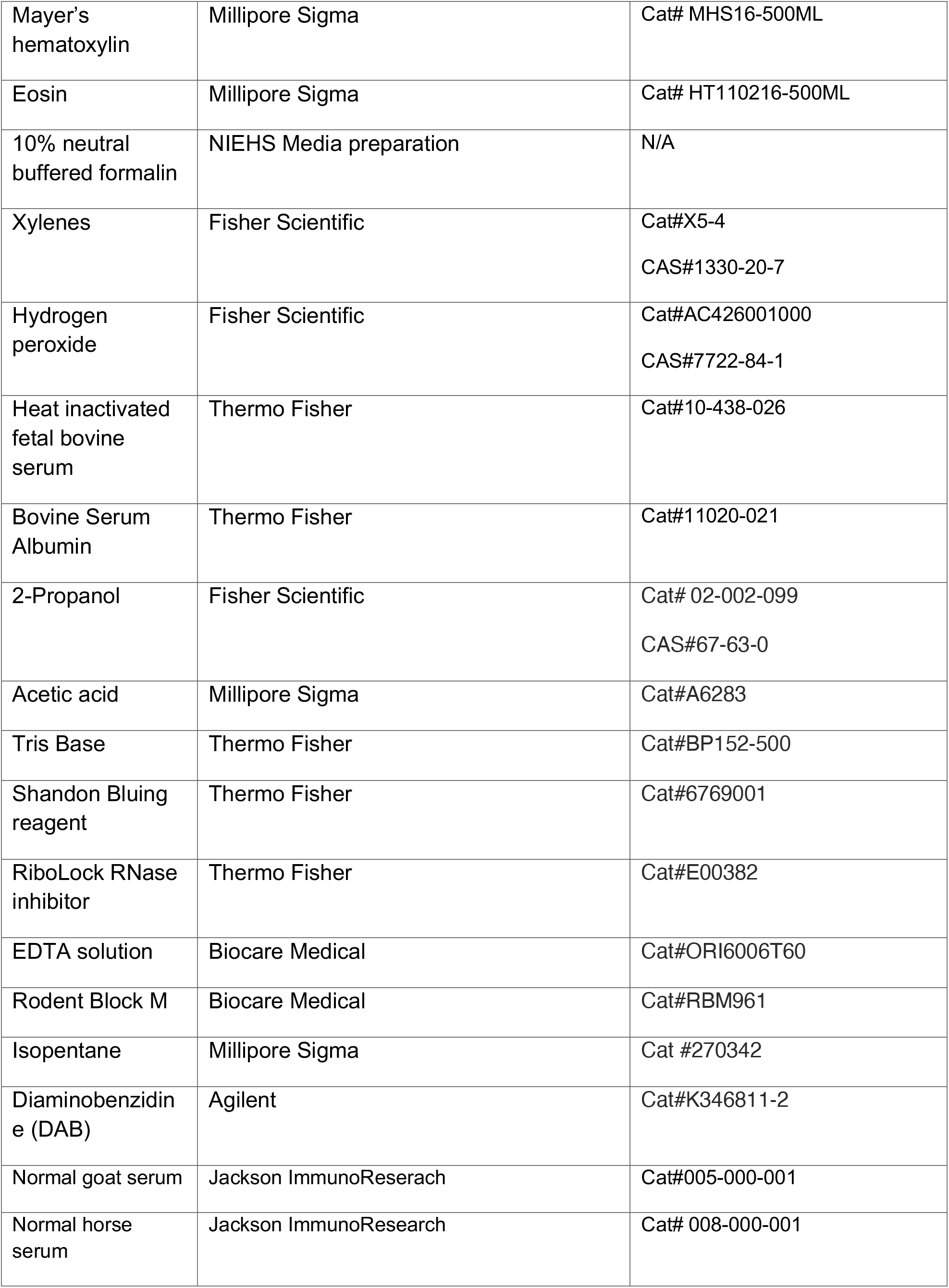
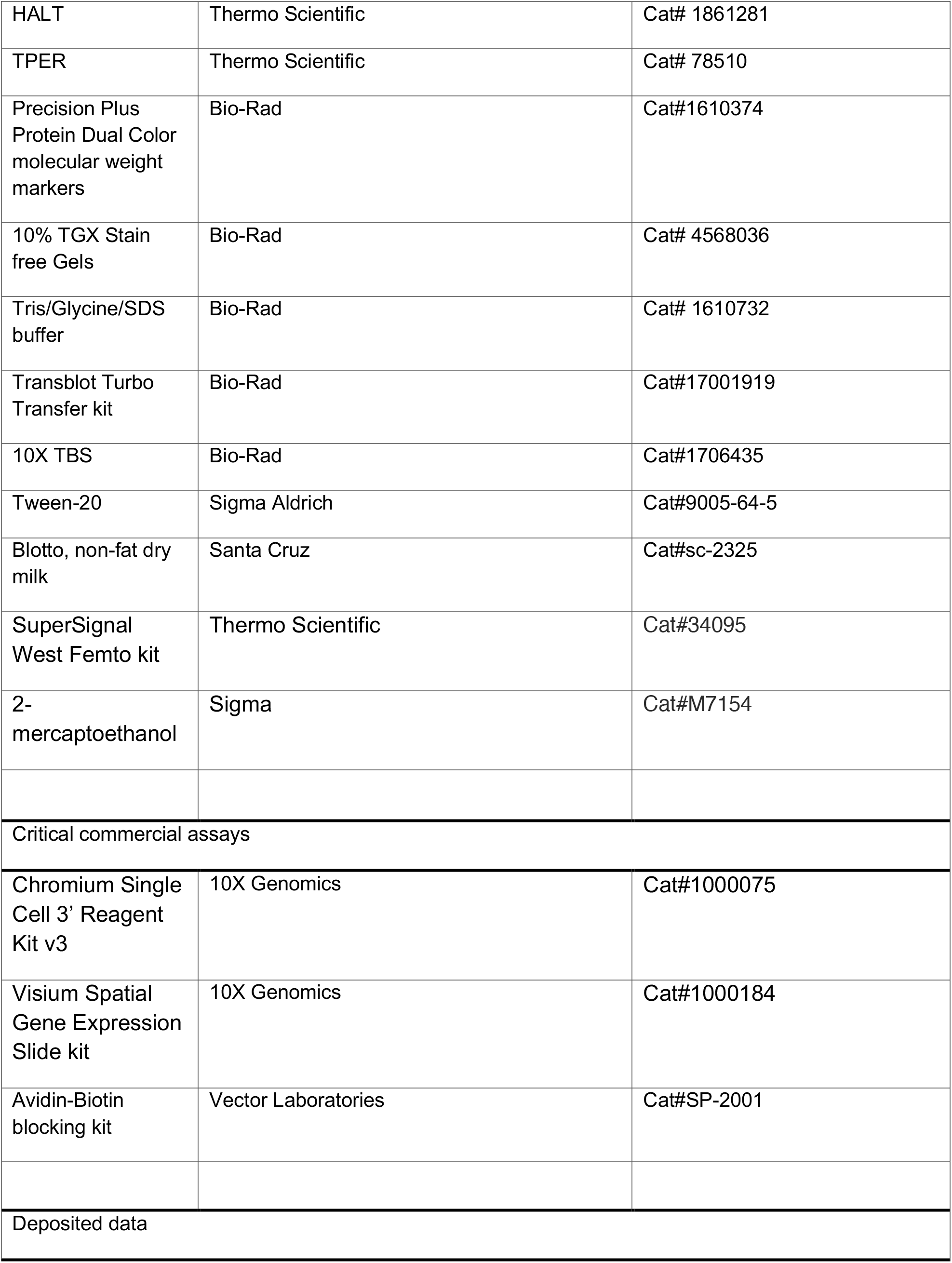
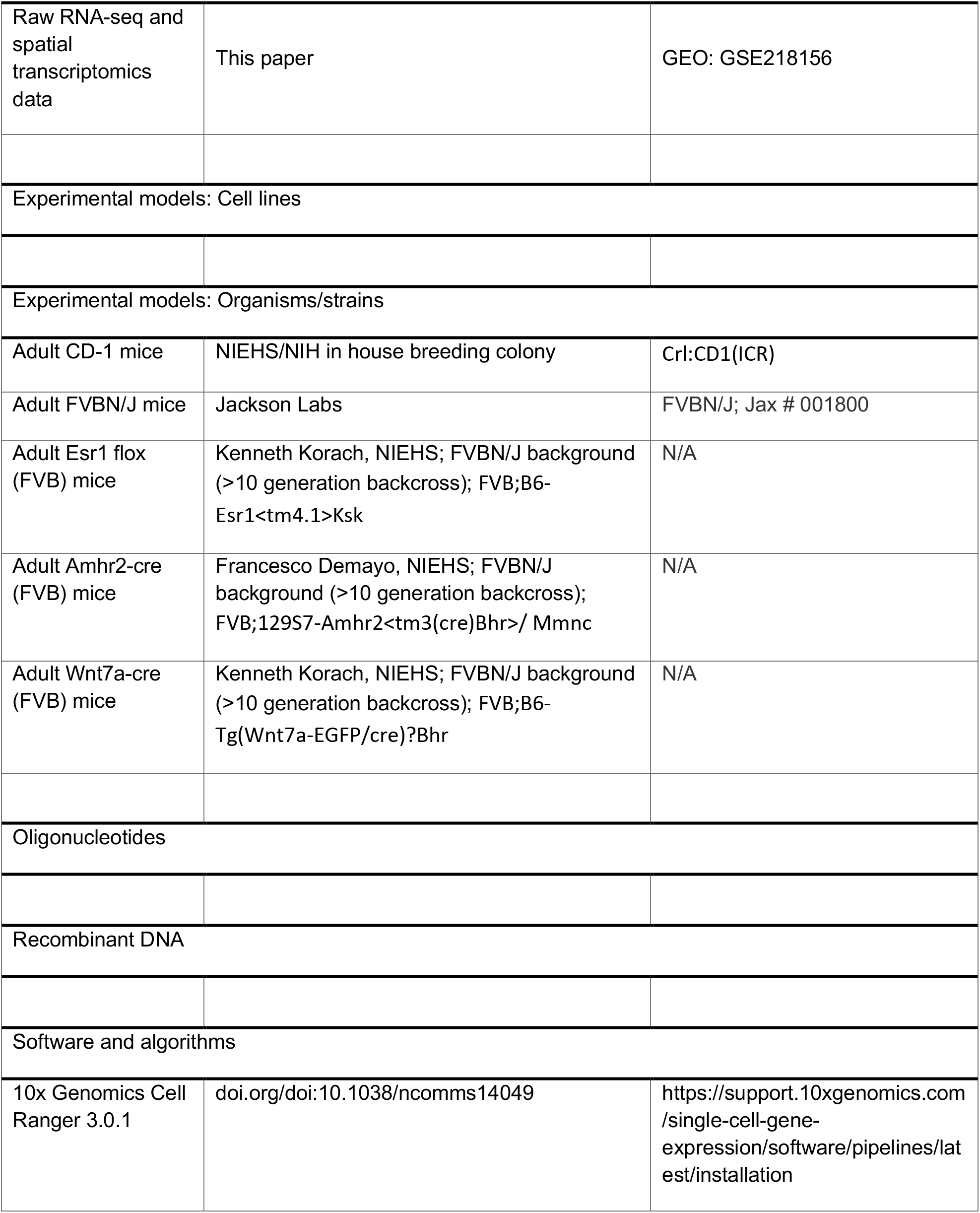
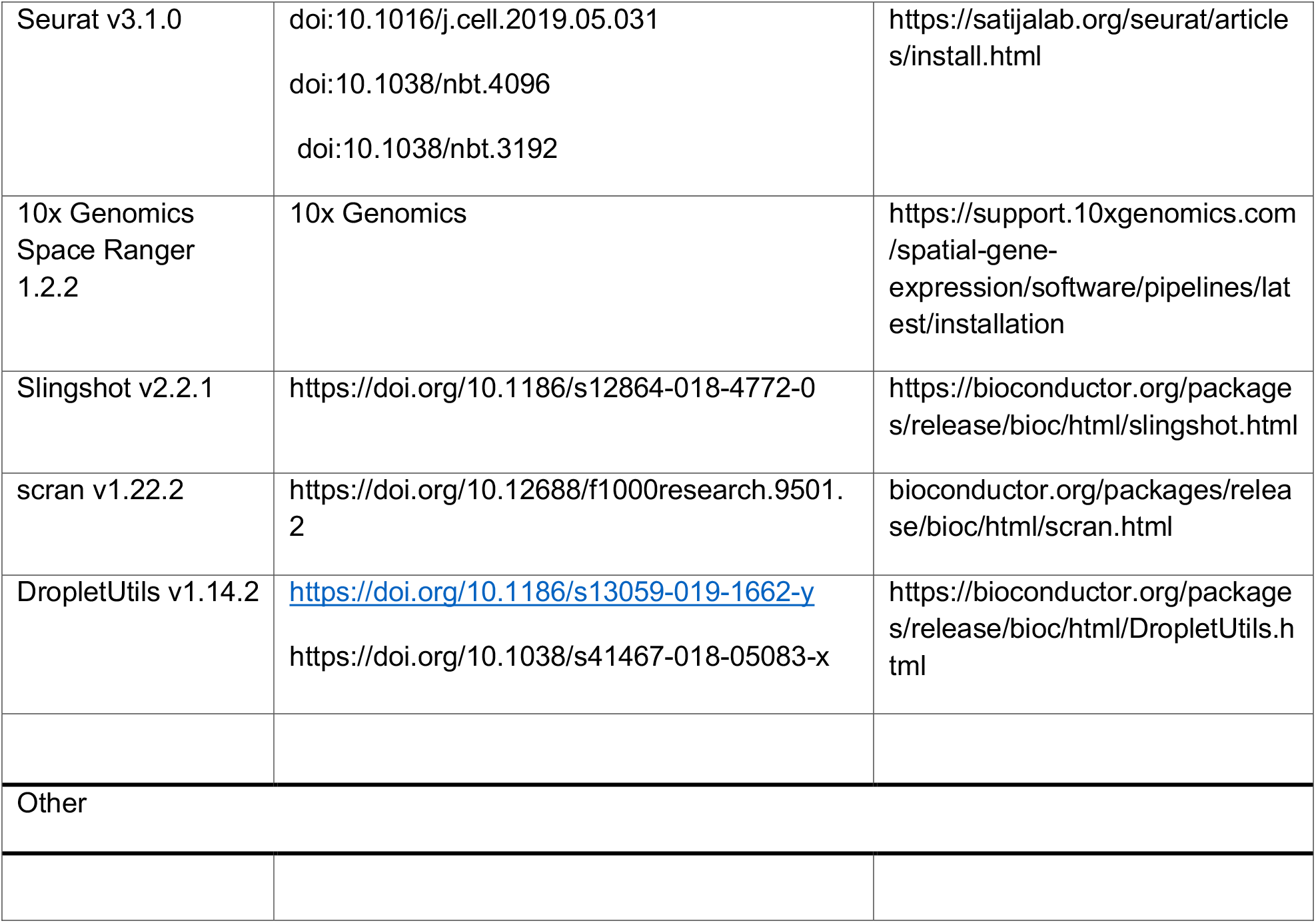

---

**Figure S1 – related to Figure 1:**
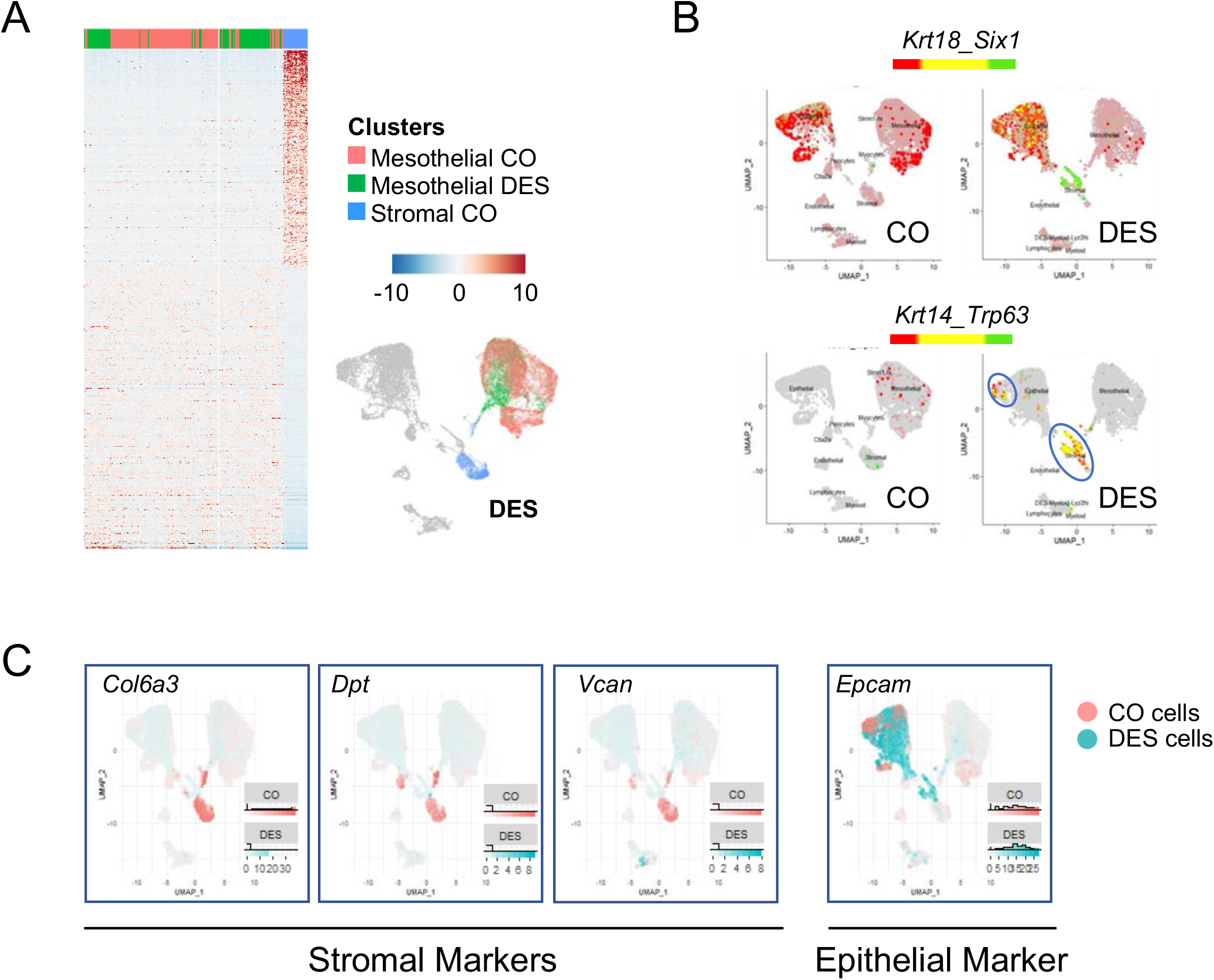
Identification of uterine cell types captured by single cell RNA-seq. A) Heat map of mesothelial and stromal cell markers in CO and DES epithelial cells. Expression is Pearson Residuals from the SCTransform method. Umap is from Figure 1A with cell types indicated by color. B) Dual feature plots of epithelial and basal cell markers (*Krt18, Krt14, Six1* and *Trp63*). CO (left) and DES (right). Colors for each gene (red or green) are indicated above the UMAPs and yellow indicates overlapping expression. C) Feature plots of stromal markers *Col6a3, Dpt* and *Vcan* and epithelial marker *Epcam* using the integrated Umap of all cells from CO (peach) and DES (teal). Expression is the same as A.

**Figure S2 – related to Figure 2:**
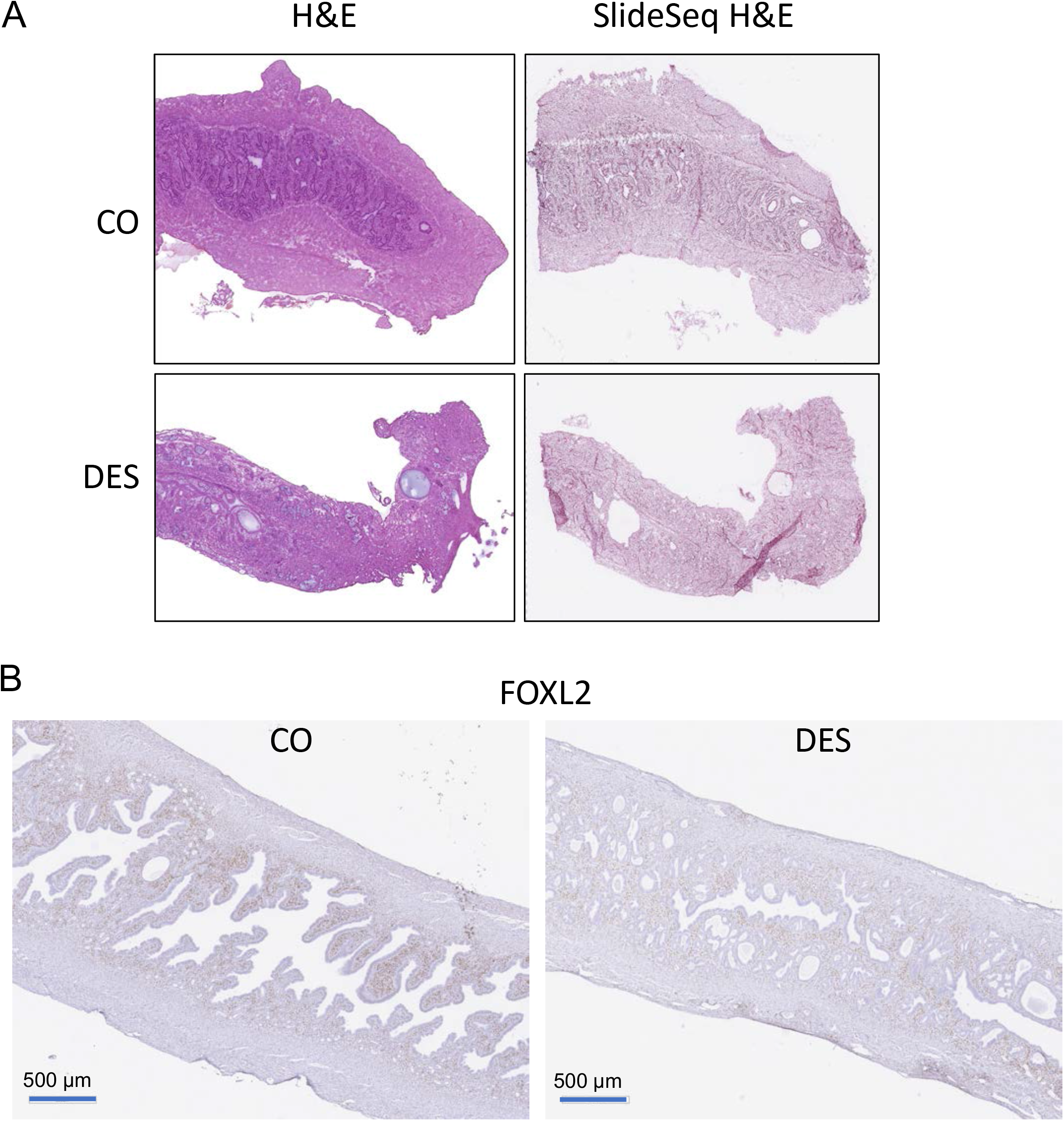
Stromal cell gene expression is altered by DES as evidenced by spatial transcriptomics. A) Uterine tissue sections from CO and DES used for spatial transcriptomics. H&E stain of tissue section adjacent to spatial transcriptomics section (left) and tissue sections used in spatial transcriptomics (right). B) FOXL2 IHC in CO and DES. Scale bars indicate size (microns=μm) on each panel.

**Figure S3 – related to Figure 3:**
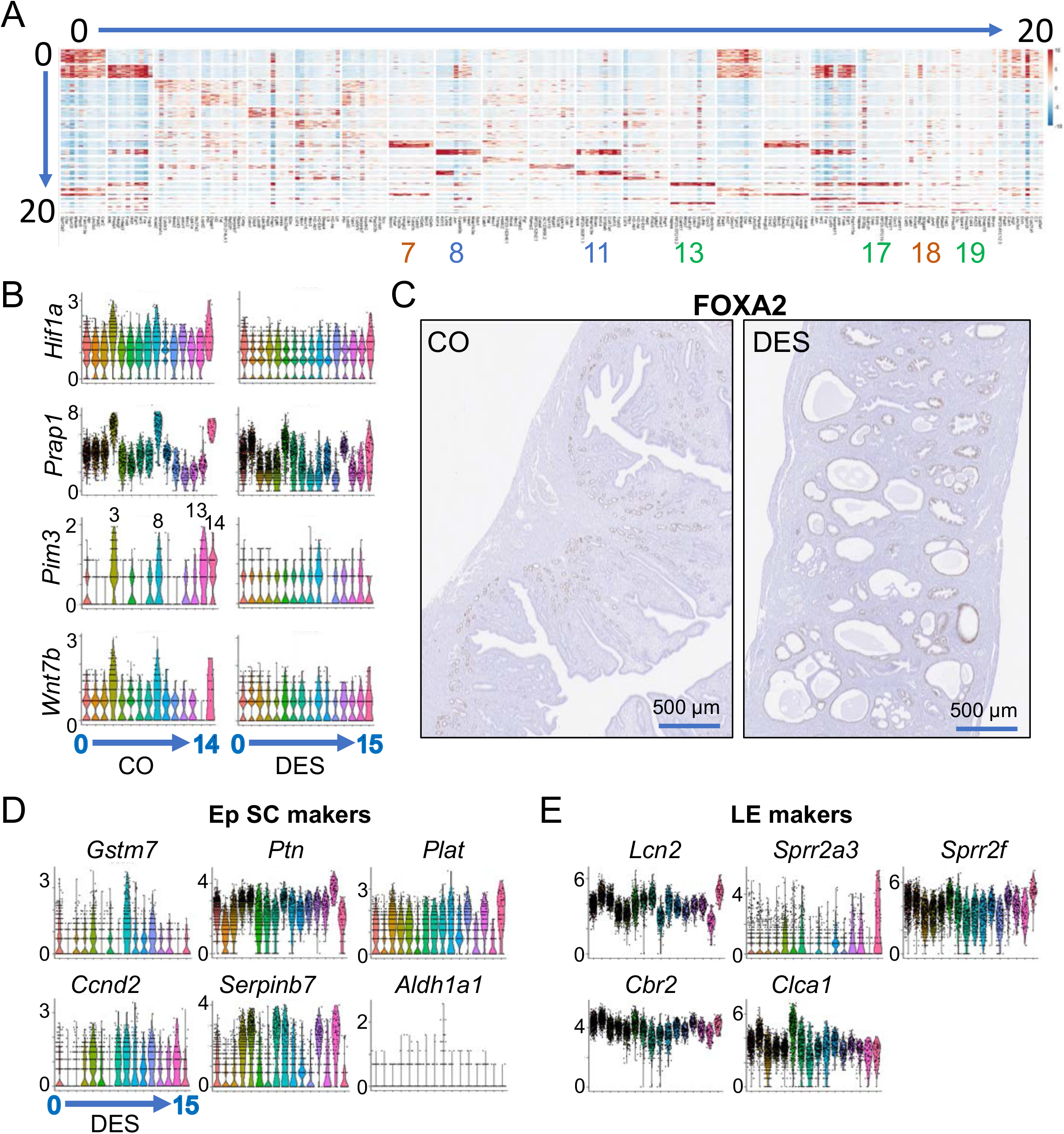
Epithelial cells from DES exposed mice lack subtype identity. A) Heat map of top DEGs of each cluster from integrated epithelial cell Umap. Expression is Pearson Residuals from the SCTransform method. Clusters are indicated across the top and select clusters indicated: Basal cells (8, 11); GE (13, 17 and 19); overlapping CO and DES clusters (7 and 18). B) Violin plots of developing GE markers (*Hif1a, Prap1, Pim3* and *Wnt7b*) for CO and DES. Cluster numbers are indicated below violin plots. Expression is natural log transformed counts and is indicated for each gene. C) FOXA2 IHC in CO and DES. Scale bars indicate size (microns=μm) on each panel. D) Violin plots of top 5 LE and Ep SC markers from Figure 3F. Cluster numbering and expression are the same as in panel B.

**Figure S4 – related to Figure 5:**
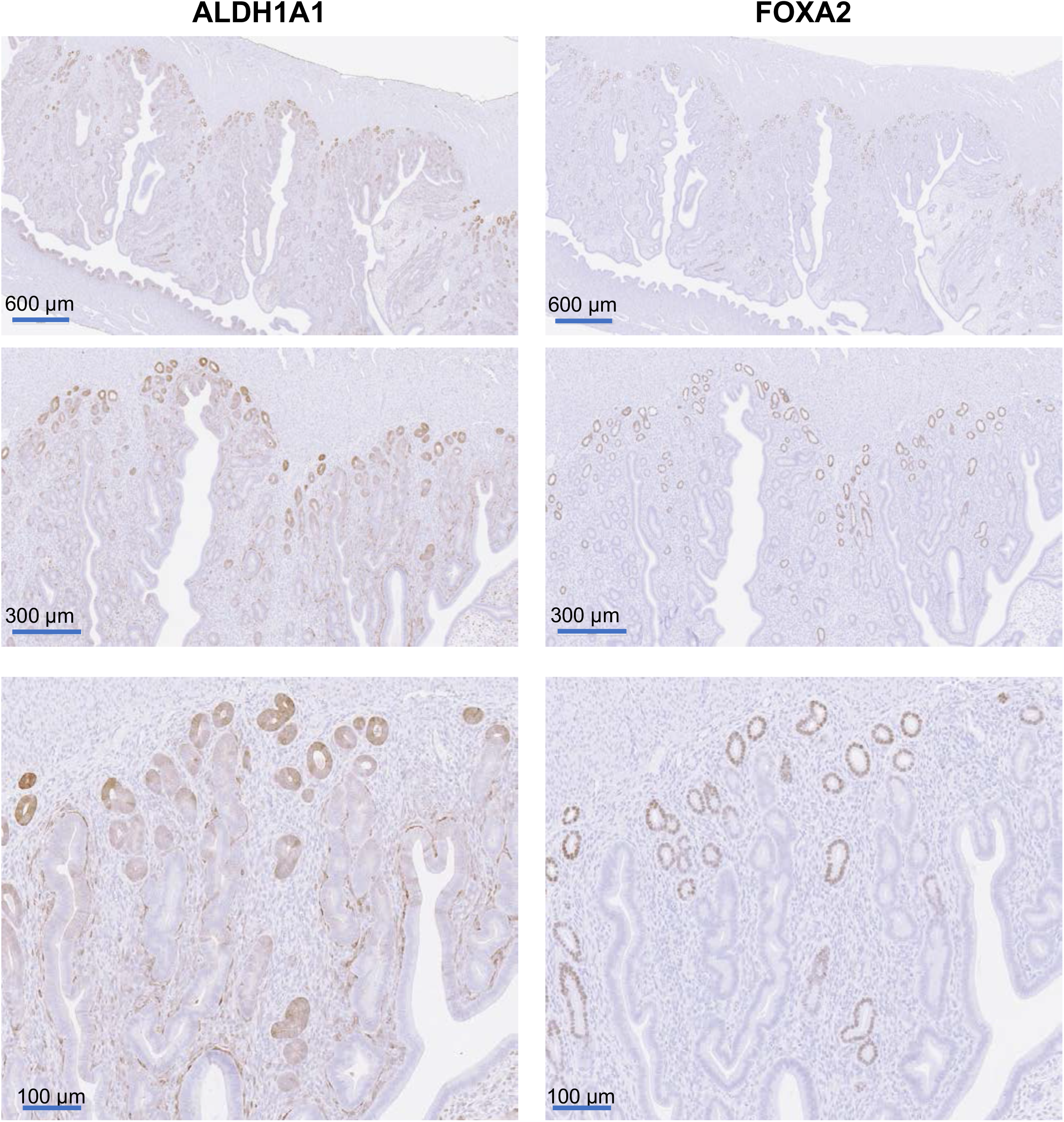
ALDH1A1 and FOXA2 expression is overlapped in CO mature GE but not in EpSC. ALDH1A1 and FOXA2 IHC in CO uteri. Adjacent sections shown at increasing magnification.

**Figure S5 – related to Figure 6:**
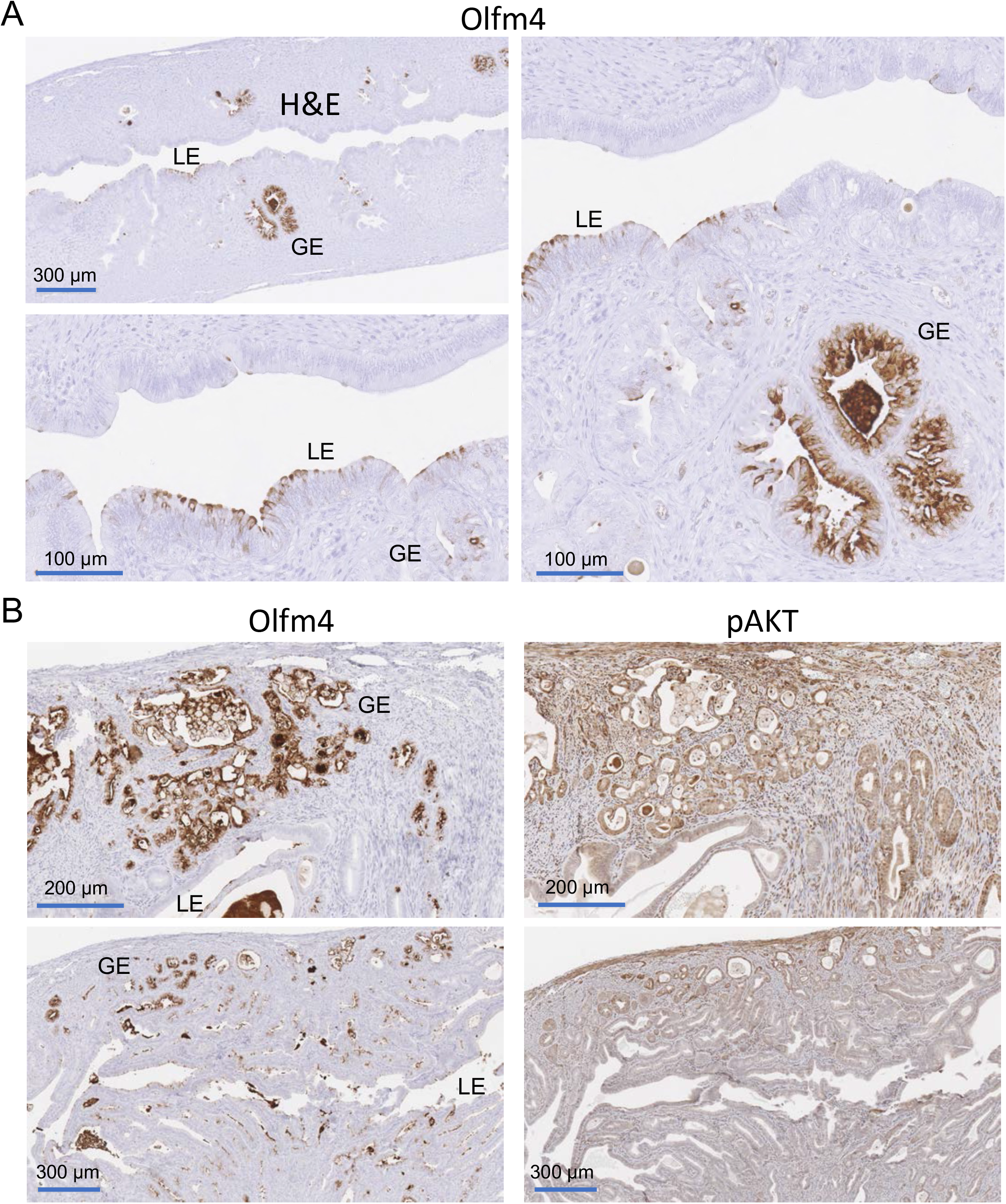
OLFM4 and pAKT are overlapped in DES-exposed uteri at 12 months of age. A) OLFM4 IHC in CD-1 12 month old DES uteri. B) OLFM4 and pAKT IHC in FVBN/J 9 month old DES uteri. Scale bars indicate size (microns=μm) on each panel. GE and LE are indicated.

